# Overcoming Biosynthetic Limitations to Enhance Bacterial Polyketide Production

**DOI:** 10.1101/2025.09.16.676551

**Authors:** Sofia C. Bravo, Jingyi Hu, Susanna Kushnir, Marius Brandenburger, Frank Schulz

## Abstract

The objective of this study was to enhance the production of monensin and its derivatives in *Streptomyces* sp. ATCC 15413. To this end the contributions of medium composition and enzyme engineering on polyketide biosynthesis were assessed. Enzyme engineering was implemented through a single-point mutation in KS5 of the polyketide synthase (PKS). This mutation increased premonensin productivity up to 29-fold, revealing and alleviating a rate-limiting step in the multi-enzyme biosynthetic pathway.

Medium optimization proved comparably effective, raising titers by at least an order of magnitude across strains. Moreover, medium optimization and ketosynthase mutagenesis acted additively in the premonensin strain, further boosting its production.

Overall, our findings show that medium optimization is the dominant factor in maximizing monensin yields, while enzyme engineering can deliver targeted benefits in specific contexts.

## Introduction

*Streptomyces* spp. are a rich source of bioactive compounds, from which a substantial share of medically relevant natural products are produced.^1,2^ Many other antimicrobial or antitumoral drugs come from natural products but have been structurally optimized to improve its pharmacodynamic and kinetic properties.^3–6^

Recent advances in the engineering of natural product biosynthetic pathways have opened the access to fully biosynthetic new-to-nature natural products.^7–12^ This could hypothetically be used for the creation of bioactive compound libraries. At its current level of development, many new compounds become detectable but they are rarely produced in meaningful quantities.^12,13^

Enhancing biosynthetic yields of natural product derivatives is therefore crucial to allow their testing and mass production. For this task, researchers have traditionally relied on strain engineering approaches, targeting either regulatory pathways to modulate flux control or, more recently, introducing targeted mutations within polyketide synthases (PKS) to boost productivity.

Still to this day, the design of productivity-enhancing mutations remains a formidable challenge, largely due to the inherent complexity of the producing organisms. Historically, this complexity has led to the widespread adoption of classical, random strain mutagenesis strategies, which, while effective, often lack precision.^14–16^ An example of strain engineering is the production of monensin, a polyether ionophore, by a specifically cultivated Bulgarian variant of *Streptomyces*, strain A519 (from now on, BP strain).^17,18^ Monensin is biosynthesized by a type I polyketide synthase (PKS) and shows a remarkable bioactivity, e.g. as an antibiotic and antiparasitic in animal husbandry and, more recently, antineoplastic activity has been described *in vivo*.^19^ The monensin PKS consists of eight large enzymes (MonAI–MonAVIII) organized into 12 modules. This modular assembly line synthesizes a linear polyketide backbone, which is subsequently modified by a set of post-PKS tailoring enzymes to yield the final bioactive compound (Figure S1).

Here, we describe an experimental platform to enhance productivity in *Streptomyces* strains towards premonensin, monensin and its new-to-nature derivatives using targeted mutagenesis and medium optimization.

We used *Streptomyces* A495 strain as a model for premonensin and its derivatives. As monensin model strains, we used Wild-Type (WT) monensin and Bulgarian Producer (BP) strains.

## Results and Discussion

### Design and generation of single point mutants

Despite decades of effort and several notable successes in engineered PKS biosynthesis, the field continues to struggle with low product yields and limited scalability.^20–22^ In a previous study from our group, we generated monensin derivatives by inactivating different reducing domains in the PKS. We obtained low titers of the new-to-nature products and proposed that one key contributor to this could be biosynthetic bottlenecks caused by inefficient processing of the new-to-nature intermediates.^13^

As first approach to increase the production of polyketides, we considered a direct mutagenesis of the monensin PKS itself ^13,23^, allowing us to circumvent the complexities of traditional metabolic engineering.

Inspection of liquid chromatography–mass spectrometry (LC-MS)-based metabolomics data of *Streptomyces strain A495* revealed irregular patterns in the biosynthetic intermediates along the PKS assembly line, specifically the signal of the intermediate from module 4 (M4) was notably higher compared to downstream intermediates.^13^

This may be attributed to a mass spectrometric detection bias, where the chemical properties of the M4 intermediate enhances its detectability.^24,25^ Varying ionization efficiencies among different compounds require rigorous calibration and the use of appropriate internal standards to ensure accurate quantification.^26–30^

Alternatively, this observation could indicate a real biosynthetic bottleneck, suggesting a rate-limiting step in the biosynthetic pathway, specifically within module 5, which is located directly downstream. If the high M4 signal indeed reflected increased abundance rather than an analytical artifact, the ketosynthase in module 5 (KS5) could be expected to possess a reduced processivity.

Enzymological studies on PKS indicate that the KS domain often constitutes the rate-limiting step in each module. This kinetic bottleneck is typically attributable to several factors: The KS substrate specificity, the positioning required for catalysis, and the conformational adjustments necessary for effective turnover relative to subsequent processing steps.^31^

To shed light into this, we examined the sequences of ketosynthase (KS) domains within the monensin PKS (Figure S2).^32^ A specific motif spanning positions 224-227 in KS5 (Figure 1A) caught our attention due to a deviation from the conserved consensus sequence observed in other modules.^13^We hypothesized that this divergence may underlie the relative low processivity of KS5 and designed an experiment to test this hypothesis.

**Figure 1:**
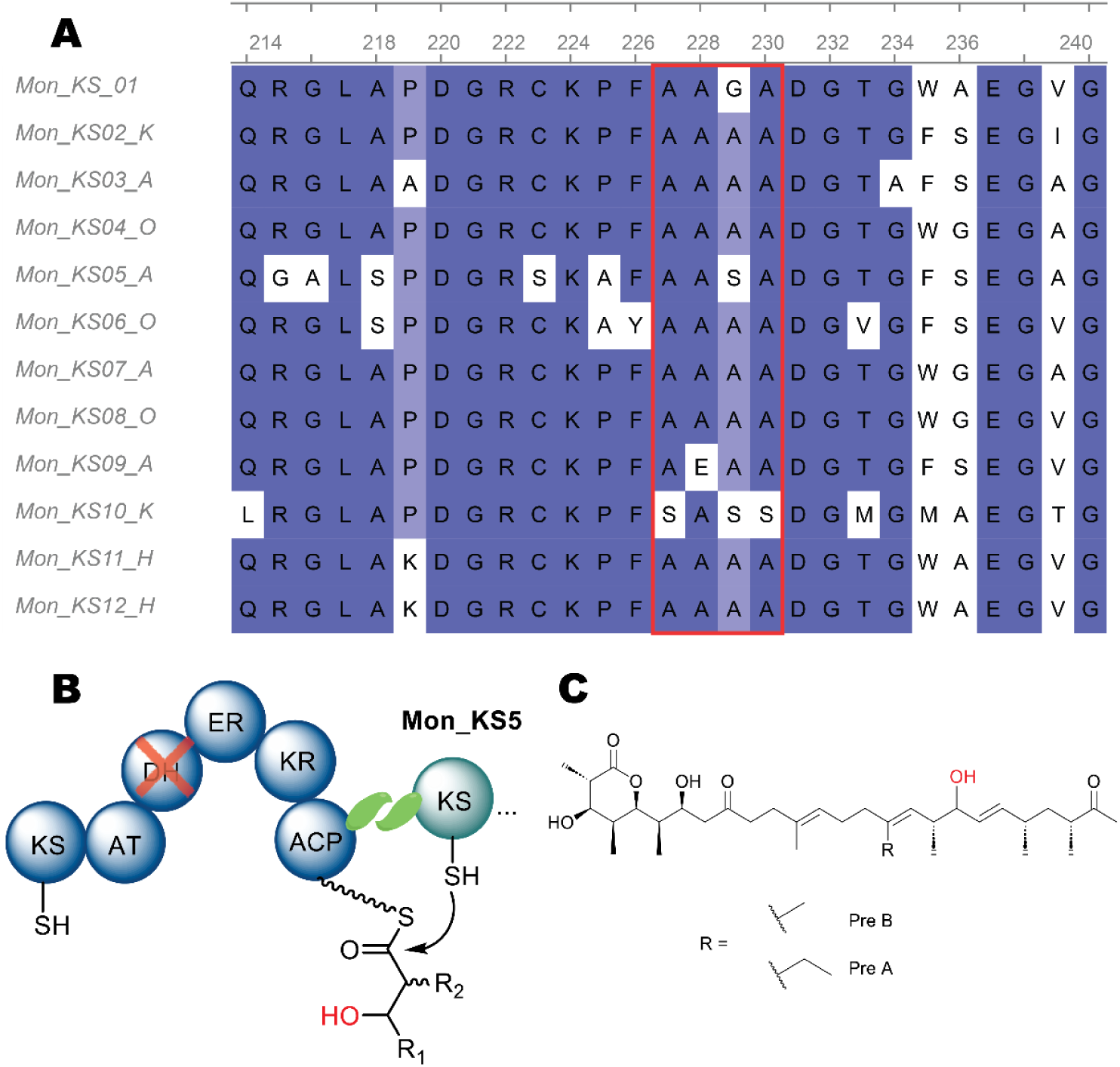
A) Excerpt of a Multiple Sequence Alignment of monensin KS domains (full alignment in supplementary information, Figure S2). The red box highlights positions 224–227. 75 % of aligned KS domains carry an alanine at position 226, whereas KS5 contains a serine instead of the consensus alanine, as does KS10. The analogous mutation in KS10 yielded no viable clones in any of the tested recipients. Multiple further sequence motifs and their effects will be discussed in a separate manuscript on the enzymology of ketosynthase domains. B) Engineered DH4^0^ variant: Inactivation of the dehydratase (DH) domain from module 4, where the catalytic histidine in the active center is mutated to phenylalanine, thereby inactivating the domain without compromising its fold. This modification slows the reductive step, leading to the accumulation of oxidized premonensin derivatives. C) Structure of DH4^0^ premonensin A and B.

We introduced the KS5 S226A mutation into two variants: the A495 variant (ATCC 15413 ΔmonCIΔmonBI/II*, vide supra*) and the DH4^0^ variant of A495; which features a more extensive modification, with both the post-PKS enzymes and the dehydratase (DH) domain in module 4 inactivated. See *SI, Tables S1 & S2* for oligos and plasmids used. This modification results in the production of a hydroxy-derivative of premonensin (Figure 1B,C). DH4^0^ was selected to investigate whether the S226A mutation would not only alleviate a potential bottleneck in the A495 variant, but also influence the yield of premonensin derivatives, indicating a more general impact on PKS activity. In a control experiment, the same mutation was introduced into *Streptomyces* wild-type (WT) and the Bulgarian Producer (BP).

### Mutation S226A effect on premonensin titers

To assess the influence of the S226A mutation on premonensin biosynthesis as an example for a new-to-nature polyketide, we evaluated three clones of the A495 reconstituted wild type (rWT) and four A495-S226A clones to validate reproducibility and inter-clonal variability. Analysis via high-resolution LC-MS (LC-HRMS) was focusing on the key intermediates and end-products of the biosynthetic pathway, premonensin A (PreA) and premonensin B (PreB).^33,34^As shown in Figure 2A and B, the S226A variant exhibited a near 2-fold increase in PreA and PreB LC-MS integrals compared to the rWT. These results were accompanied by abundance changes in shunt products (Figure 2A). Specifically, the S226A mutation led to a significant decrease in M4 and a concomitant increase in M5B, M6A/B, M7A, and M11A/B. The results corroborated our initial hypothesis, indicating that the replacement of a potentially suboptimal amino acid with its consensus counterpart can indeed enhance overall productivity, thereby suggesting that a single, targeted mutation may be sufficient to alleviate a primary yield limitation. To control the LC-MS-results on the S226A mutation with an orthogonal experimental approach, quantitative Nuclear Magnetic Resonance (qNMR) spectroscopy was used, based on its previously demonstrated reliability for precise absolute quantification of premonensin.^40^ Although qNMR requires a more labor-intensive workflow with reduced throughput and sensitivity compared to LC-MS, its application provides an intrinsically quantitative readout.^41–46^

**Figure 2:**
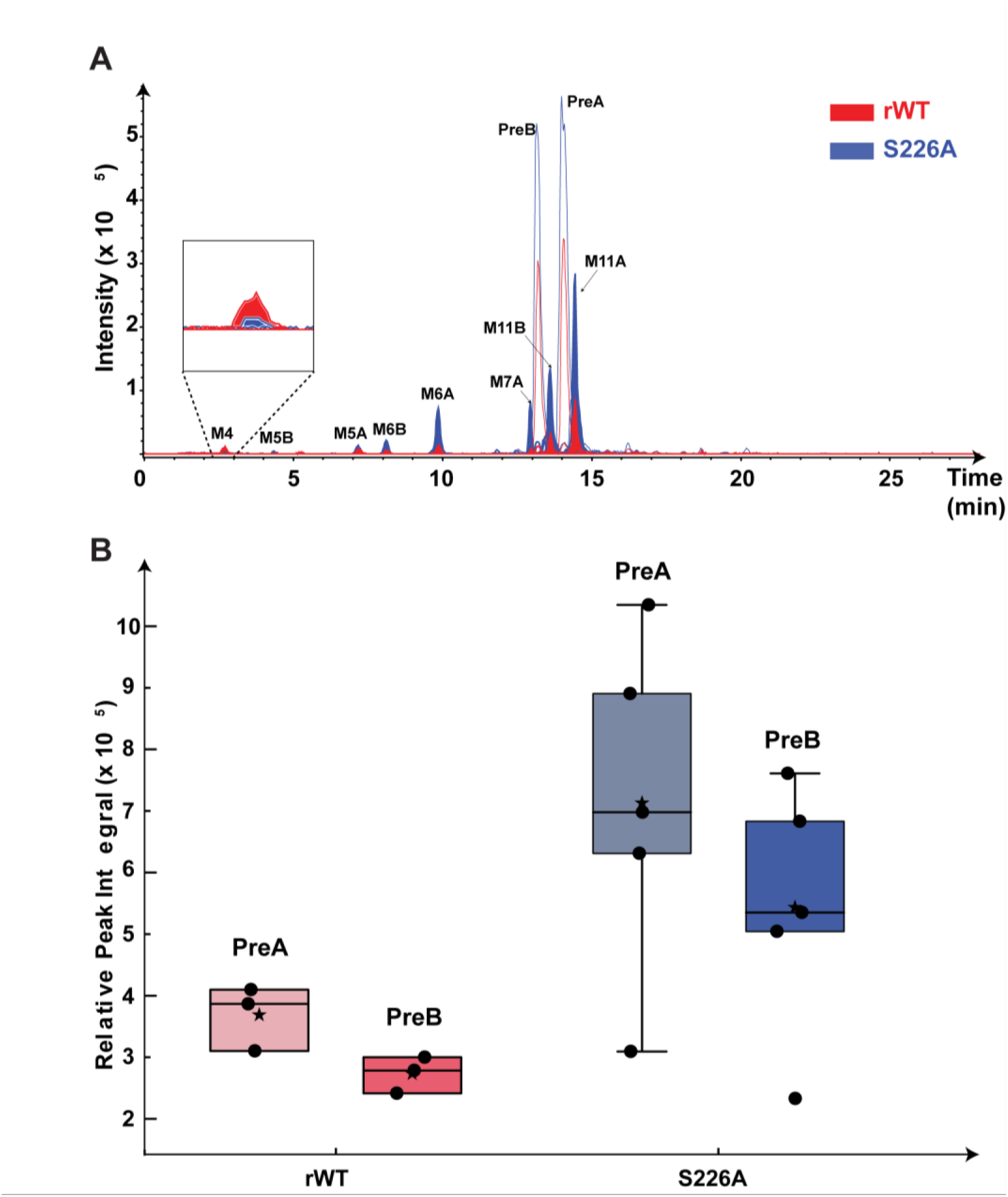
A) Intermediate and product profiling extracted ion chromatograms ([M+Na/H/K/NH_4_]^+^) of intermediates (M) and final products PreA and PreB, revealed substantial changes in variant S226A (blue) compared to the reconstituted wild type (red). There was an increased abundance of intermediates M5B, M6A/B, M7A, M11A/B, and final product PreA/B; as well as a decreased abundance of intermediate M4. B) Boxplot of comparative analysis of PreA and PreB production. Normalized LC-MS peak integrals for PreA and PreB in A495_rWT and A495_S226A (mutant) variant are shown with black dots. The line shows the median and the star shows the average normalized signal. There is a significant impact of the S226A mutation on increasing product yields. A list of spectra processing steps, all adducts of interest, as well as their fragmentation patterns can be inspected in the SI, Section C) HPLC-MS Analysis.

To this end, multiple consecutive fermentation batches were used, each involving a direct comparison between rWT and the S226A variant. Clones with different LC-MS product integrals were deliberately selected to evaluate the effect across a wider range of putative clone productivities (See Figure 3C). In Figure 3A the whole NMR spectrum of an enriched premonensin extract is shown. The olefinic proton (in green) was used for the quantification, correlating to the signal of the internal standard (in pink). The Figure 3 B and C show that the qNMR data support the trend observed by LC-MS, but reveal an increase by one order-of-magnitude in premonensin A and B productivity attributable to the S226A mutation. This substantial enhancement provides robust validation of our hypothesis, confirming that the KS5 domain is indeed a limiting factor in the overall premonensin PKS assembly line. This result strengthens the hypothesis that substituting a suboptimal amino acid with the consensus counterpart can enhance overall biosynthetic productivity.

**Figure 3:**
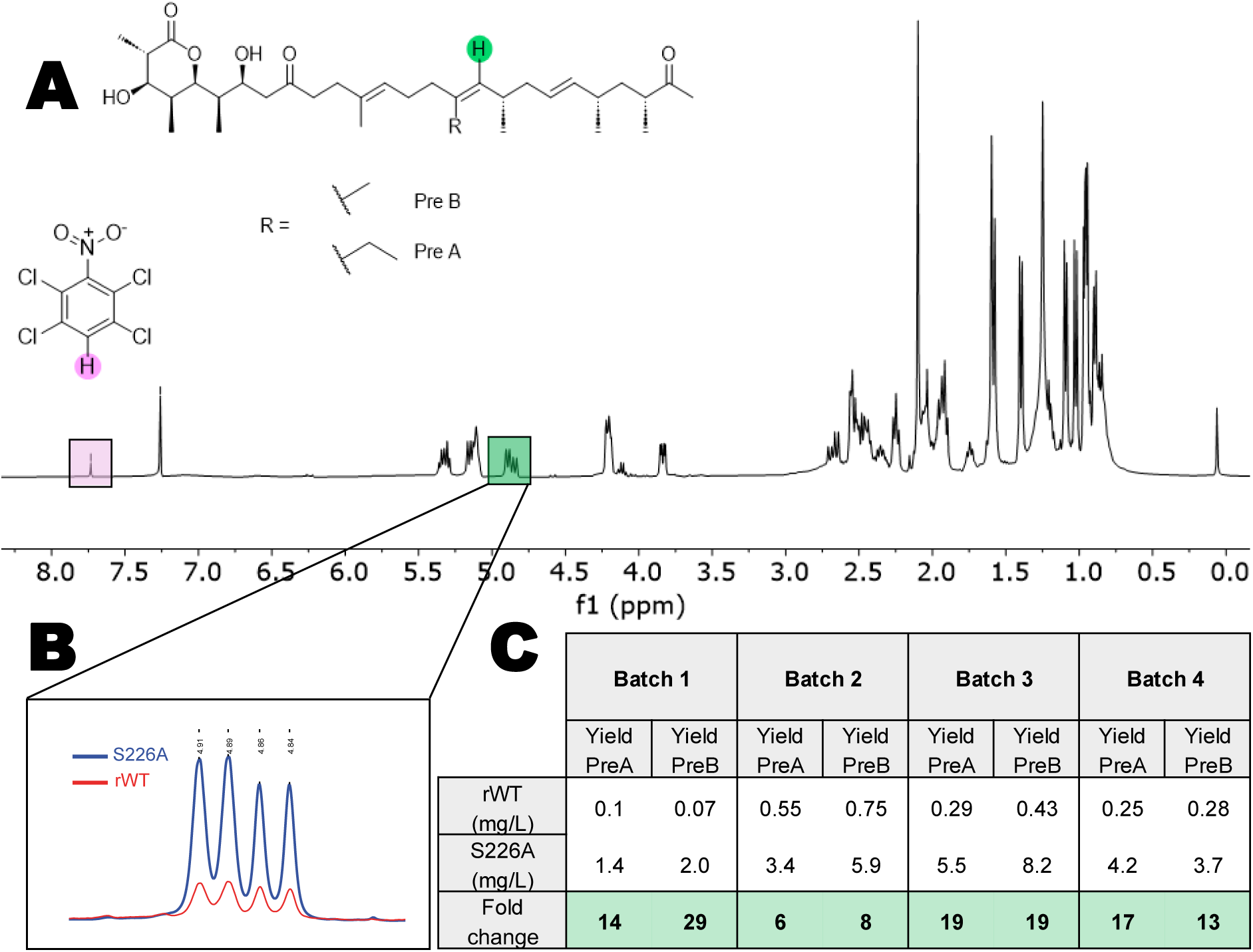
A) 1H-NMR spectrum for PreA and PreB of an A495_S226A variant sample, highlighting PreA (ethyl group at C17) and PreB (methyl group at C17). The characteristic olefinic proton is highlighted in green. Absolute quantification of PreA and PreB was achieved using 1,2,4,5-tetrachloronitrobenzene as an internal standard (singlet at 7.74 ppm, pink) in a known concentration. B) Magnified view of the NMR signals used for quantifying PreA and PreB in: A495-rWT (red) and A495-S226A (blue) PreB: 4.90 ppm, PreA: 4.85 ppm. C) Absolute quantities of PreA and PreB across variants and batches, providing an overview of the production levels. The results show that the LC-MS measurements were reliably showing a trend but underestimated its effect. Each batch was carried out with different clones, apart from batch 2 which was also supplemented with more XAD16 resin for in situ product adsorption.

### Mutation S226A effect on the premonensin derivatives titers

Following the investigation of the S226A effect on the strain A495-rWT, we extended our analysis to include nine clones of each the low-producing variants DH4^0-^rWT and DH4^0^-S226A. These variants carry a deactivated DH domain, in which the catalytic histidine in the active center of the domains is replaced by a functionally inert phenylalanine side chain. This lowers the activity of the DH domain and leads to the production of DH4^0^ premonensin A and B (Figure 4A). ^47^ Due to the inherent lability of the resulting allylic alcohol in this compound, the major products remain PreA and PreB.

**Figure 4:**
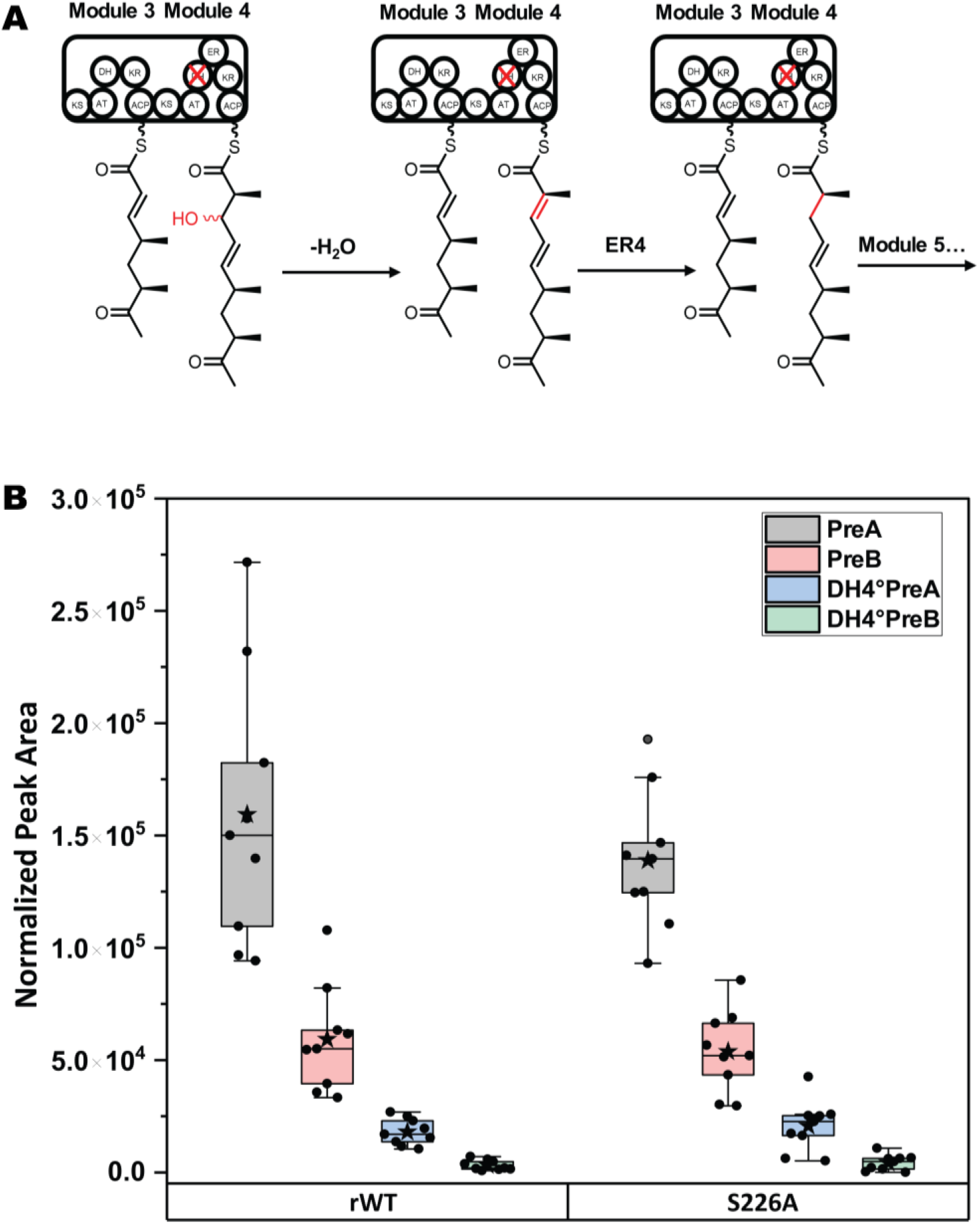
Effect of mutation S226A on the DH4^0^ strain. A) Intermediates from modules 3 and 4 are shown, with the inactivated DH4 domain marked (red X). The oxidized M4-DH4^0^ intermediate is to some extent excluded from further processing, presumably due to substrate-specificity of KS5. Instead, spontaneous water elimination from the allylic alcohol intermediate can be anticipated, facilitated by conjugation with the adjacent C=C bond and thioester, yielding the intermediate shown. This step likely represents the rate-limiting event in the biosynthetic cascade, unaffected by downstream KS domain mutations. After its formation, the olefinic intermediate undergoes reduction by the ER4 domain like the intermediate in the native module, before being passed to module 5 with some delay. B) Comparative quantification of biosynthetic intermediates. Normalized LC-MS peak integrals for PreA, PreB, DH4^0^PreA, and DH4^0^PreB are compared between DH4^()^-rWT and DH4^0^-S226A variants.

These clones were analyzed by LC-MS using the same method as for premonensin. In contrast to its effect on the A495 variants, the S226A mutation in this context does not significantly improve the production of PreA, PreB, or the hydroxyderivative (Figure 4B).

This result suggests that the main bottleneck in the biosynthetic pathway of this variant is the acceptance of the hydroxylated new-to-nature intermediate from module 4, which is not remedied by mutation S226A.

In light of all of the above, it can be concluded that this mutation does neither affect the extender unit nor the nascent polyketide specificity of the KS domain but instead its innate processivity.

### Structural Insights of S226A Mutation

Next, to investigate the underlying mechanism of the observed increase, we used AlphaFold 3 to predict the structures of the S226A variant and wild-type KS5, employing the sequences ACP4-^C-terminal^docking domain4 and ^N-terminal^docking domain5-KS5_S226A/WT as templates (see SI, section D).^47^ The WT model revealed a KS-active site channel architecture similar to that proposed for pikromycin PKS, featuring a side entrance for substrate transfer from ACP4 to KS5 and a bottom entrance for downstream catalysis (Figure 5A).^48^ Residue S226 is located on the surface of the KS domain, near the side entrance of the binding channel, in close proximity to the modeled phosphopantetheine (pPant)-attached Ser46 (DSL motif) of ACP4 (Figure 5B).

**Figure 5:**
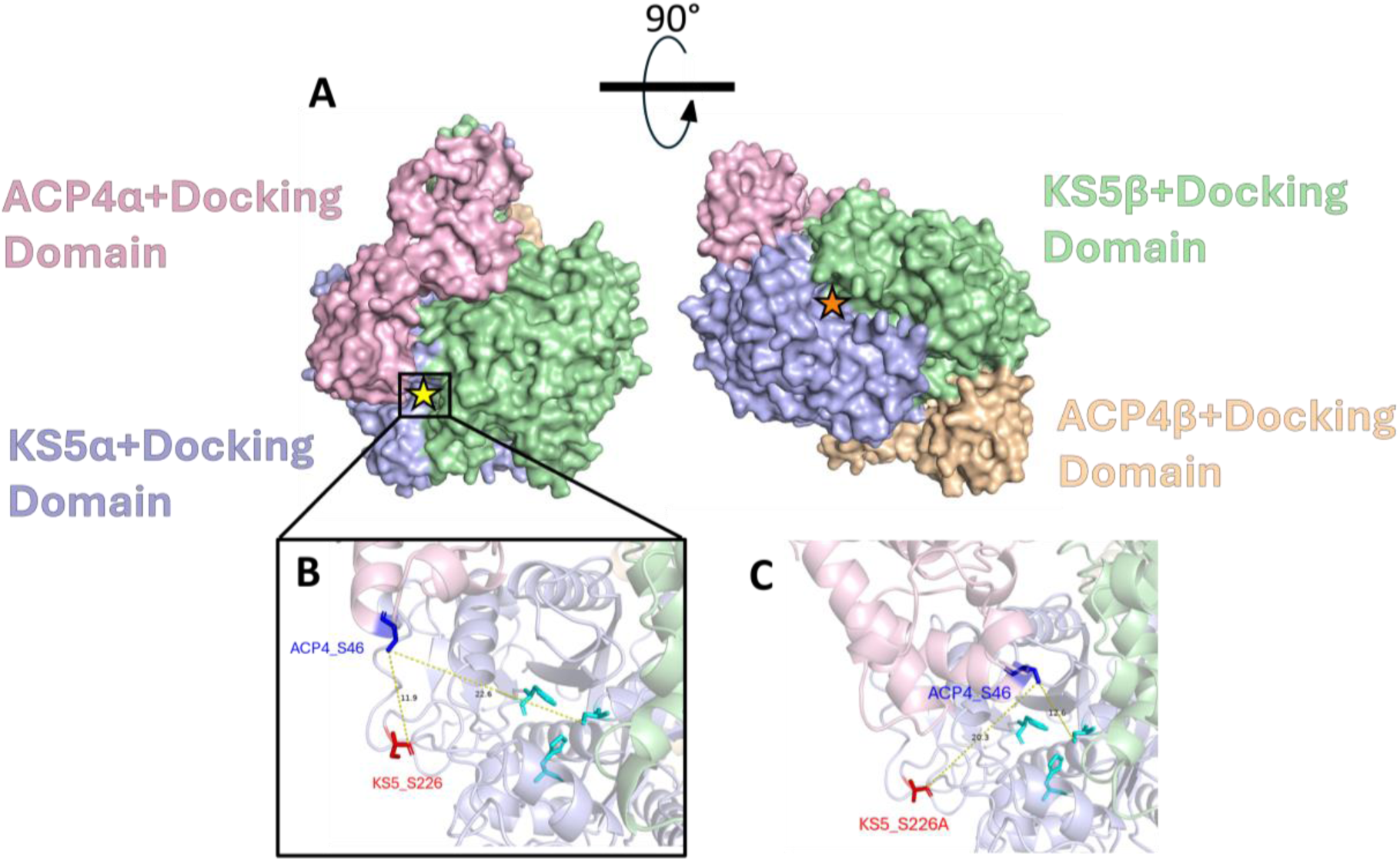
Predicted interaction between ACP4-^C^docking domain4 and ^N^docking domain5-KS5 homodimers of rWT variant and S226A variant. A) The Alphafold 3 predicted structure shows the ACP4-^C^docking domain4 homodimer (light pink and wheat) interacting with the ^N^docking domain5-KS5 homodimer (light blue and green). The side entrance of the KS5 active site is marked by a yellow star, and the bottom entrance is indicated by an orange star. B) A magnified view of the side entrance in the predicted wild-type structure shows that the key Ser46 residue (dark blue) in ACP4 is positioned in proximity to both the mutated residue Ser226 (red) and the catalytic Cys169 (cyan). The catalytic triad His-His-Cys in the KS5 domain is shown in cyan. C) In the predicted S226A variant structure, Ser46 in ACP4 appears closer to Cys169 and slightly more distant from Ala226 compared to the wild type, suggesting an altered relative positioning of ACP4 and KS5 though the structural basis for this shift remains unclear.

In the predicted S226A variant structure, the ACP4-KS5 relative positioning is altered with ACP4-Ser46 now closer to KS5-Cys169 and more distant to KS5-A226 (Figure 5C). The mutation may enhance enzymatic productivity by modulating the interaction between ACP4 and KS5. Specifically, the predicted structures suggest that substitution of serine with alanine enlarges a hydrophobic surface patch, potentially increasing the affinity between ACP4 and KS5 during the chain translocation step.^49^ Although S226 has not been directly confirmed as part of the ACP–KS interface, structural alignment with resolved KS domains places it within the region corresponding to the proposed ACP docking surface, suggesting a possible role in mediating or stabilizing this interaction.^50–52^ While the structural model provides a useful starting point, the interaction confidence remains modest, and further studies will be needed to clarify the precise mechanistic impact of this residue.

### Effect of Mutation S226A on monensin titers

Mutation S226A was able to increase premonensin titers in the mutated strain. The question arose whether the bottleneck in the biosynthesis of premonensin would also be the main bottleneck in the entire biosynthesis of monensin, including its post-PKS machinery. To investigate this, mutation S226A was introduced into monensin producer WT and BP strains, and the resulting variants were grown using fermentation conditions as used for premonensin (SM16) and with the BPM regime (commonly used for the industrial strain).^17^

As can be seen in Figure 6, the mutation influences the WT grown in SM16 medium, leading to a more than 2-fold increase of the median monensin yield. In the BP strain, the mutation effect is not statistically significant.

**Figure 6:**
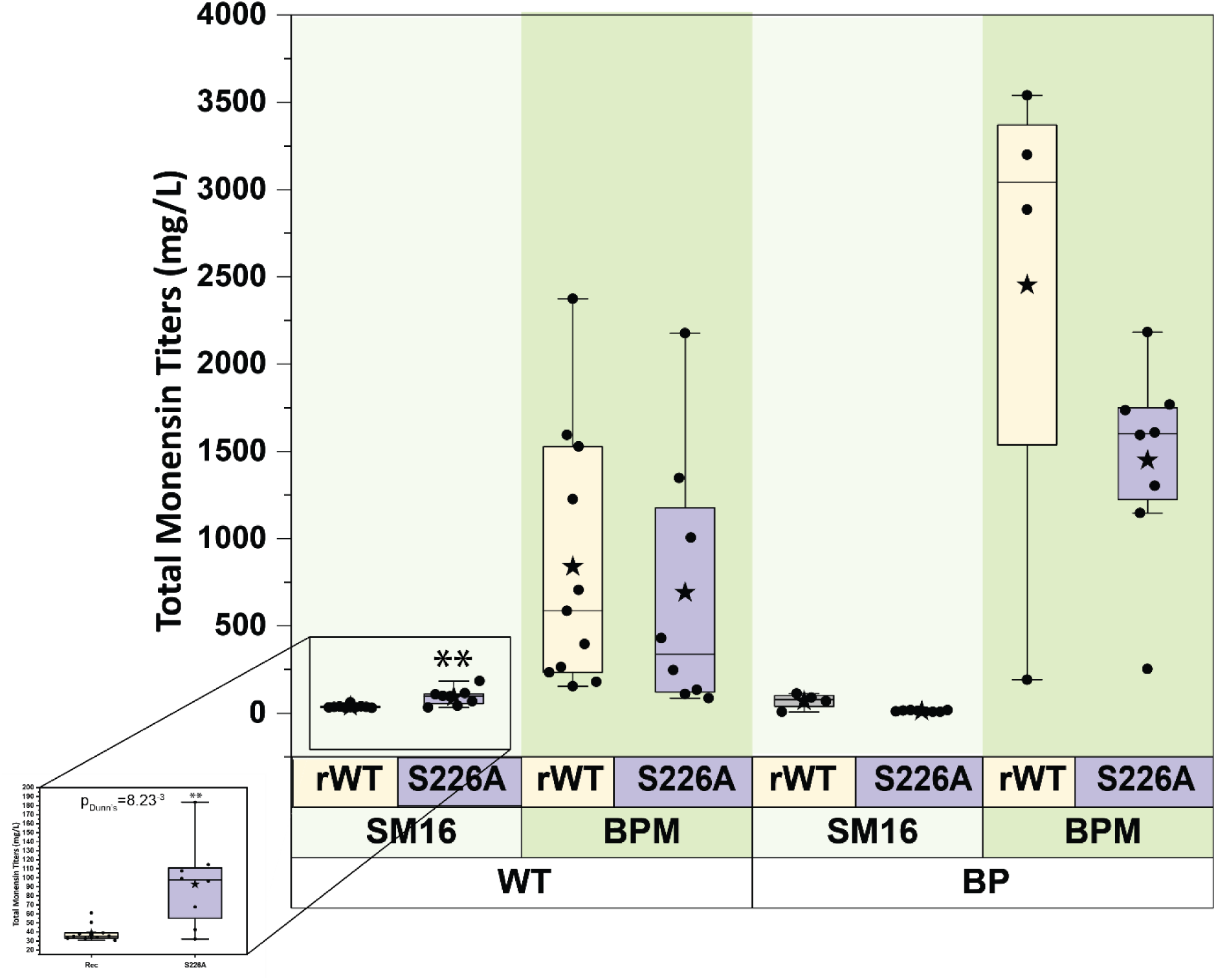
Effect of mutation S226A on WT (left) and industrial (BP, right) monensin strains. Light green is used for SM16 and darker green for BPM media. Reconstituted WT strain is shown in light yellow and the mutated strain on purple. Biological replicates are shown as black dots, the median as a line and the average measurement as a star. Non-parametric tests (Dunn’s, Origin 2024b version) were applied between each pair of rWT vs S226A. The S226A mutation only significantly increased titers in rWT grown in SM16 medium, with a p-value of 8.23^-3^. The increase in the media of the titers is 2.79-fold. WT titers with SM16 media are zoomed in for easier inspection. See SI Section E for more information about the quantification method.

The mutation had overall no effect when any of the two recipients was grown in BPM medium. This indicates that KS5 acts as a bottleneck in SM16 medium, regulating global monensin production. However, in complex medium, this step no longer appears to be limiting.

Any effect of the mutation in these experiments is strongly overridden by the statistical difference between monensin production in the minimal medium (SM16) and in the rich medium (BPM).

### Media Optimization in Monensin Production

The observation of a strong medium effect in the characterization of the S226A variant (Figure 6) prompted us to deeper assess the impact of media composition on productivity.

Initially, single colonies from both strains were tested to evaluate differences in production titers (Figure 7 A). Mycelium grown on SFM agar served as the starting point, with standardized plugs used as inoculum for subsequent fermentations.

**Figure 7:**
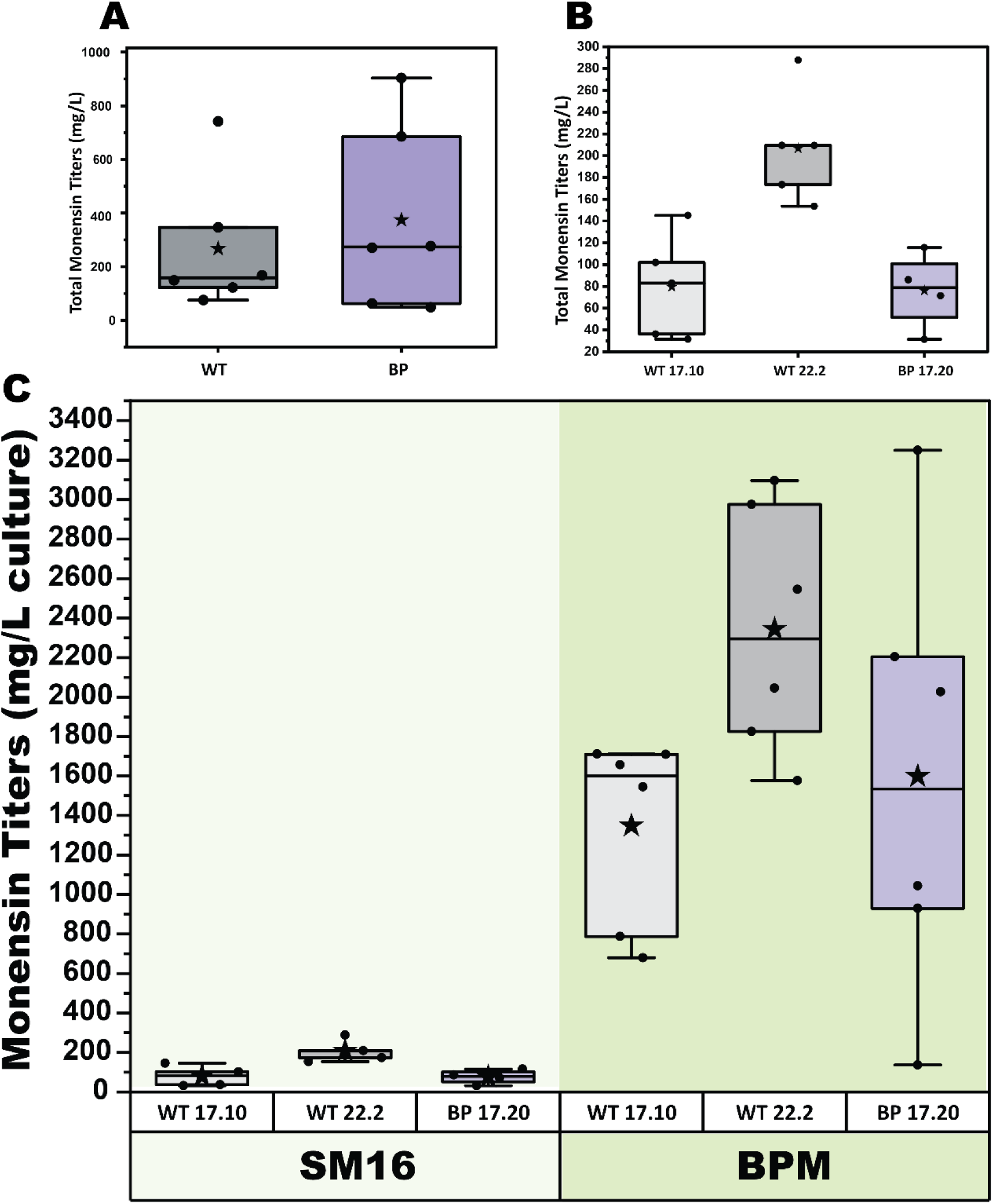
Effect of medium on monensin production in WT and industrial (BP) strains. A) WT and BP spore stocks were diluted, and single colonies were picked and grown separately to test monensin titers in the BPM regime. Boxplots show individual samples as dots. The WT distribution shows that most clones produce lower titers than the BP strain. See SI Figure S14, for clones’ individual titers. B) Monensin titers in SM16 medium. Clone WT22.2 produces significantly more than its counterparts (p-values from Dunn’s test: 0.0305 vs. WT17.10 and 0.02934 vs. BP clone, Origin 2024b). This panel is a zoomed-in version of the larger comparison shown in C. C) Comparison of titers across clones in SM16 and BPM media. Biological replicates are shown as black dots (each value is the average of two technical duplicates). The median is indicated by a line and the mean by a star.

We screened over 100 individual colonies over several generations and found substantial clone-to-clone variability. We decided to closer inspect the highest-producing WT clone (WT 22.2), a second tier WT clone (WT 17.10), and the best producing clone from the BP strain (BP 17.20) (see Figure 7B).

These 3 clones were grown in six biological replicates and their monensin productivity in the two distinct media regimes was evaluated: the defined medium SM16 and the optimized Bulgarian production medium (BPM) protocol, specifically tailored for the industrial strain. Cell dry weight (CDW) analysis of 5-day-old SM16 cultures showed comparable cellular densities for both strains (see SI, Figure S15). CDW measurements in BPM or Bulgarian Germination Medium (BGM) cultures are inherently unreliable due to insoluble solids and oil components interfering with weight measurements and therefore were not used.

A comparative analysis of monensin production in SM16 and BPM media across these clones (Figure 7C) showed that monensin titers in BPM increased media ten to thirty-fold over those in SM16.

This result underscores that media optimization plays a greater role in monensin production than genetic variability between strains.^53–62^ The positive effect of BPM on monensin production can be attributed to multiple factors, including the provision of energy and carbon sources, as well as a soybean flour-based substrate providing a solid support for mycelial growth. Aeration, agitation, and shear stress are also crucial factors influencing *Streptomyces* growth and monensin production. By highlighting the critical role of media composition, these findings emphasize the need for tailored medium development to unlock the full productive potential of a given strain. ^63–71^

### Exploring Medium Variations for Enhanced Production

The addition of vegetable oil to an aqueous medium is a critical factor in enhancing polyketide titers, as observed in various species.^72–74^ Vegetable oil addition to bacterial fermentations is proposed to divert metabolic flux from de novo fatty acid synthesis by supplying exogenous fatty acids that feedback inhibit the native pathway, thereby increasing the intracellular availability of malonyl-CoA for polyketide biosynthesis.^75^ Soybean oil addition seems to correlate with increased titer of polyketide production in another *Streptomyces* species by upregulating the transcription of exporter genes.^76^ Specifically, it was shown that oil plays a role in achieving high productivity in BPM towards the fermentation of monensin.^17^

We substituted soybean oil with alternative vegetable oils in BPM fermentations. We hypothesized that oils with different compositions might exert distinct effects on monensin production.^77,78^

The impact of substituting soybean oil with other options was not statistically significant, although a slight trend towards increased production was observed with peanut and olive oils (Figure 8). The flexibility of the BPM fermentation medium is notable, given its ability to utilize various vegetable oils despite their chemical differences. This adaptability is especially valuable for industrial scalability under fluctuating raw material costs.^79,80^

**Figure 8:**
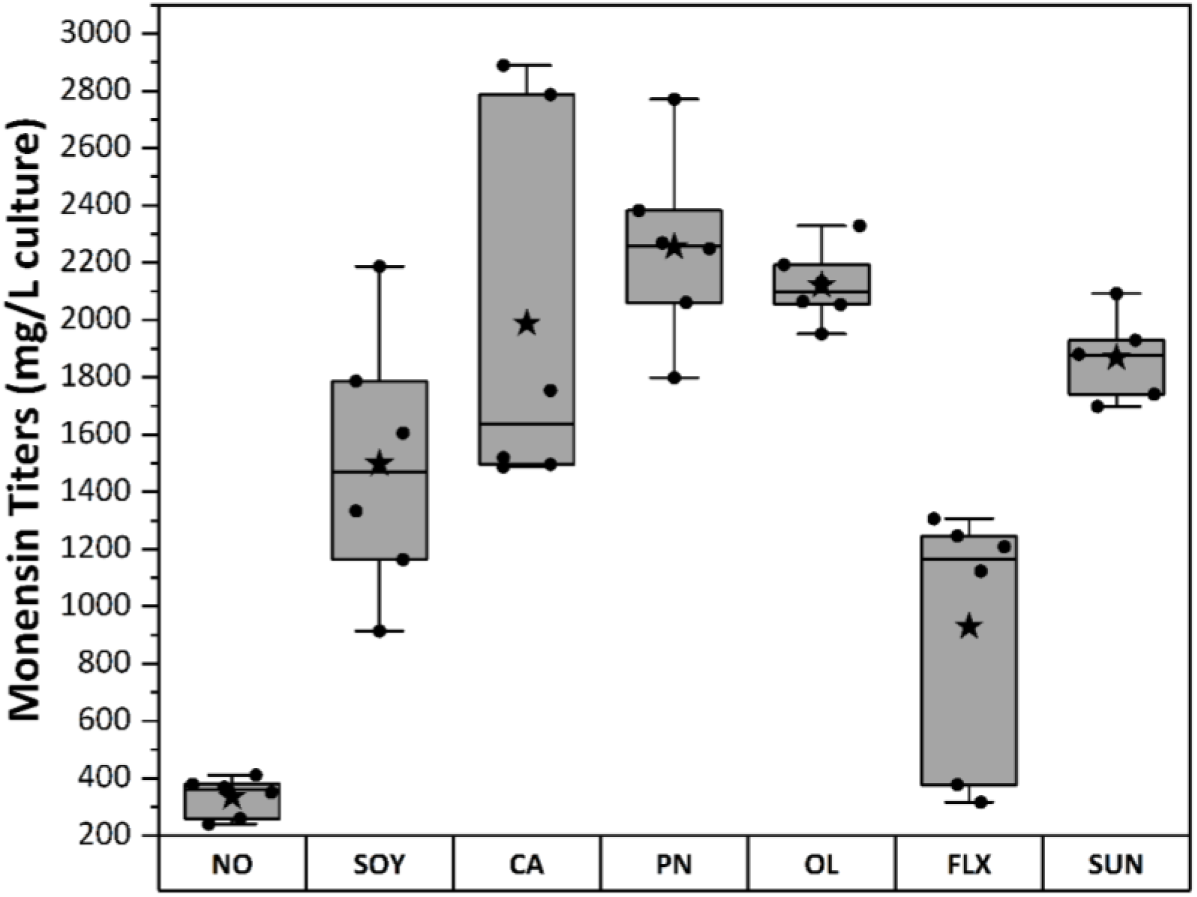
Impact of Vegetable Oils on Monensin Production in Feeding Experiments. Comparison of monensin production yields with various vegetable oils in feeding experiments. The tested oils include: No Oil (NO), Soybean oil (SOY), Canola oil (CA), Peanut oil (PN), Olive oil (OL), and Sunflower oil (SUN). Each oil was evaluated across 6 biological replicates, with 2 technical duplicates per replicate, represented by black dots. Statistical metrics are indicated as follows: median (horizontal line), mean of technical duplicates (black star). Mean rank differences are shown in SI, Figure S16.

In addition to oil substitution, we explored the potential benefits of adding reducing agents, such as thiourea or vitamin C, as well as surfactants and emulsifiers.^81,82^ Although BPM already contains a low concentration of vitamin C, we examined the effects of substantially higher concentrations. Elevated vitamin C levels increased monensin production by 80 % compared to the standard BPM (Figure S17), while thiourea was ineffective (in the concentration tested). This suggests that a better protection against oxidative stress under the high-oxygen concentration in an oil-rich medium is beneficial for the bacteria (or provides antioxidant activity for the unsaturated fatty acids in the autoclaving).^83^

We also investigated the impact of polyethylene glycol (PEG 8000) and xanthan gum (XG) on monensin production. Following sonication-assisted solubilization, two concentrations of each emulsifier were tested in both SM16 and BPM media. The results showed that in SM16, 5 g/L PEG yielded elevated monensin levels, whereas XG had no substantial impact (Figure S17&18). In contrast, BPM, with its inhomogeneous composition of solid particles and oil in an aqueous medium, showed higher emulsifier effects, with increasing XG concentrations correlating with enhanced production.

Overall, the BPM medium shows a remarkable flexibility for optimization, and we found a substantial influence of emulsifiers and antioxidants on monensin productivity.

**Beyond the norm: Testing the Medium Effect on premonensin** Following the strong positive effect of medium on monensin production, we returned to the premonensin strain to test whether the optimized BPM regime could also enhance premonensin productivity, which in our laboratory typically reaches only milligram-per-liter levels when fermentation is carried out in SM16 medium.^35,40,84,85^ Cultures of the rWT *Streptomyces* A495 strain were grown in parallel under the BPM and SM16 regimes.

Quantifying premonensin with LC-MS or qNMR as done so far was prevented by their incompatibility to the high oil content in BPM. Residual oil is intrinsically incompatible with an LC-MS system, while the high unsaturation of the vegetable oils was preventing an isolated signal for premonensin in NMR. Therefore, an alternative assay was needed, eventually resulting in the use of semi-quantitative TLC.

The observed effects are consistent with additivity, as the combination of the mutation with optimized medium conditions resulted in a detectable increase in premonensin titers (see Figure 9A). This additive relationship indicates that the mutation and medium composition exert their effects through distinct mechanisms, allowing for a predictable and cumulative enhancement of premonensin production.

**Figure 9:**
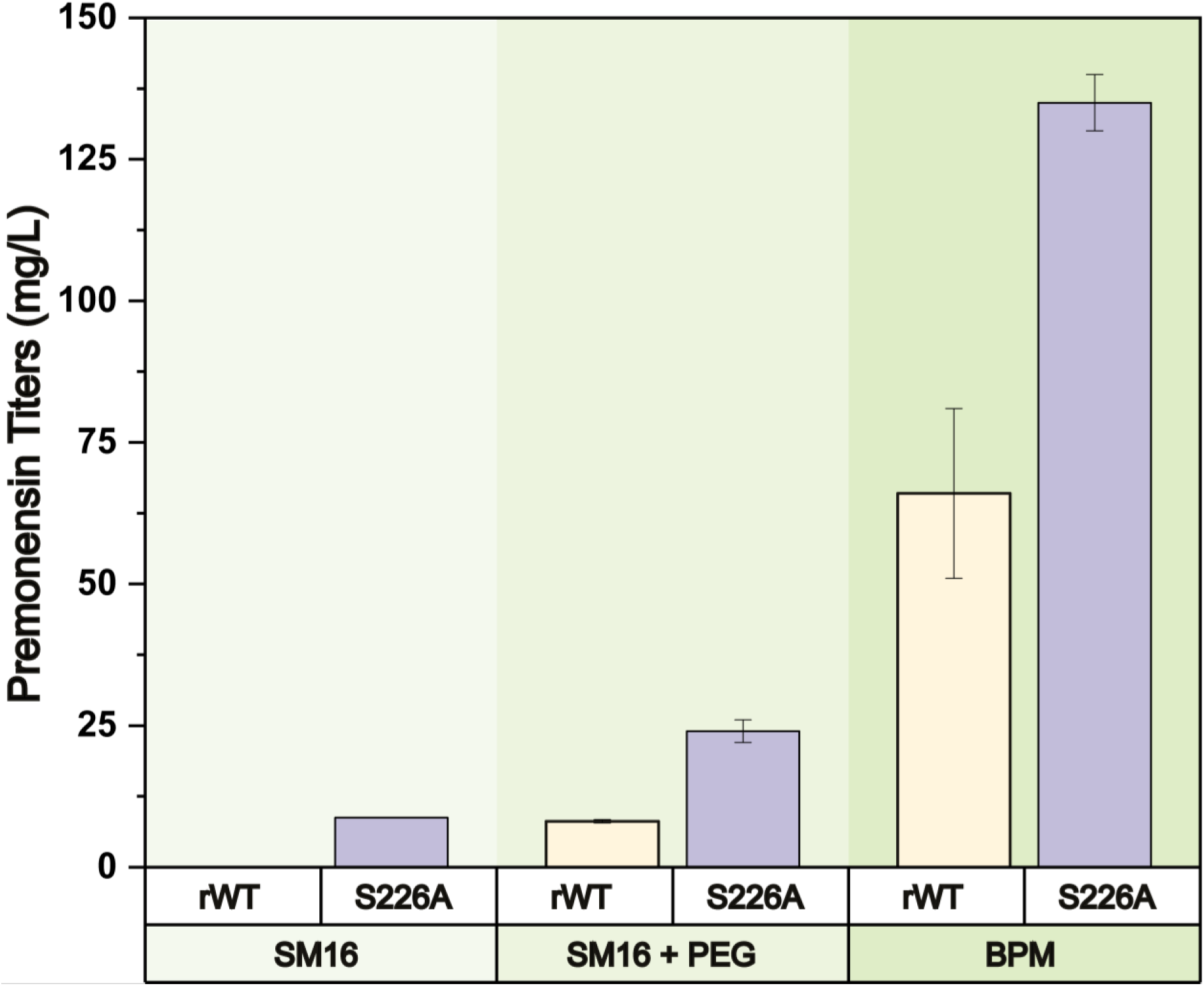
Effect of medium on premonensin production in rWT and S226A strains. Premonensin titers were calculated using densitometry of TLC plates with a premonensin external calibration curve. Each TLC plate contained all three media conditions of each biological replicate and four premonensin standard solutions. The error bars correspond to the standard deviation of two biological replicates of each condition. In rWT, premonensin is undetectable in SM16 in this assay, but faintly visible with PEG supplementation (SM16+PEG). All step-by-step processing settings are detailed in the SI2.

Future work will explore scale-up strategies and downstream processing improvements.

## Conclusion

In this study, we investigated site-directed point mutagenesis as a strategy to improve polyketide production. We found that one mutation in ketosynthase 5 of the monensin PKS enhanced productivity in premonensin. This effect likely stems from enhanced KS–ACP domain interactions, as suggested by AlphaFold modelling and is plausibly strain- and context-dependent.

By contrast, medium optimization proved to be a broadly effective strategy, increasing titers by an order of magnitude regardless of strain or mutation. This approach is likely generalizable to other polyketide producing *Streptomyces* species, with only minor adjustments to media composition. Simple additives such as antioxidants, thickening agents, or emulsifiers can be readily incorporated to further boost secondary metabolite yields, consistent with previous reports showing benefits for both polyketides and other natural products. The strong effect of medium optimization likely arises not only from providing biosynthetic precursors and energy, but also from regulatory effects such as upregulation of exporter genes in oil-rich environments.

Importantly, media optimization and targeted mutagenesis acted additively in our premonensin strain. This is particularly relevant for the generation of new-to-nature compounds, where additional bottlenecks or reduced PKS processivity often cause substantial decreases in titers compared to native polyketides.

Our findings suggest that activity regulation in PKS involves not only individual non-consensus residues but also accumulated mutations that appear to balance PKS processivity with post-PKS modifications. In the monensin producer, post-PKS processing emerged as the major rate-limiting step.

## Materials and methods

All protocols, media compositions, and instrumentation details are available in the Supporting Information (SI Section I).

### KS Mutagenesis

The S226A construct was generated by overlap-extension PCR and cloned into the pKC1139-based vector carrying a temperature-sensitive replicon using the GC-SLIC method.^86^ The GC-SLIC method involves a two-step process of PCR amplification and DNA assembly, allowing for precise mutagenesis and cloning of GC-rich polyketide synthase genes. Resulting plasmids were integrated into the chromosome by homologous recombination via conjugation. The mutant allele was confirmed by the sequencing of PCR product from total DNA.

### Liquid Cultures

Before all fermentations, tryptic soy broth (TSB) precultures were prepared from either agar plugs or frozen mycelium stocks. SM16 and BPM fermentations were performed at 30 °C with different cultivation times, with BPM involving additional germination and oil-feeding steps. The fermentation conditions, including culture volume, bioreactor type, and duration, are described in detail in the SI.

### Polyketide Extraction

Monensin, premonensin, and intermediates were extracted from cultures using ethyl acetate. For premonensin and intermediates, resin and cell paste were harvested, lyophilized, and extracted with ethyl acetate. The organic phase was concentrated and reconstituted in acetonitrile for LC-MS analysis. For quantitative NMR (qNMR) analysis of premonensin, cell/resin paste was freeze-dried, extracted with ethyl acetate under bead beating, and enriched by silica gel filtration. Monensin was extracted directly from BPM cultures by multiple ethyl acetate extractions, and combined organic phases were concentrated to yield crude extracts.

### HPLC-MS Method

An Ultimate 3000 HPLC System equipped with ISIS NUCLEODUR® C18 column (MACHEREY-NAGEL GmbH & Co. KG, 1.8 μm particle size, 2 mm diameter, 150 mm length) and a Compact mass spectrometer (BRUKER DALTONIK GmbH, Life Sciences, Bremen, Germany) with a standard electrospray ionization (ESI) was used. The mobile phase consisted of acetonitrile with 0.1 % formic acid (solvent B) and water with 0.1 % formic acid (solvent A). All solvents were of LC-MS grade. All original raw data LC-MS files used in this paper can be accessed through MassIVE MSV000099201 (doi:10.25345/C59W09C0X). HPLC and MS methods, as well as spectra processing are fully described in the SI.

### Quantitative NMR analysis

The raw products were reconstituted in 0.5 mL CDCl₃ and spiked with 0.1 mL of an internal standard solution of 1,2,4,5-tetrachloronitrobenzene (5 mg/mL in CDCl₃). NMR measurements were carried out on a 400 MHz spectrometer (Bruker Avance III HD 400 with a BBFO-ATM z-degree 5 mm sample head). To suppress carbon satellite signals, 13C decoupling was employed, and a 30 second relaxation delay was used between excitation pulses to allow full proton relaxation. Spectral processing was performed using MestReNova software. Manual phase correction was applied, and baseline correction was conducted using a third-order polynomial fit. All original raw qNMR files used in this paper can be accessed through Zenodo: https://doi.org/10.5281/zenodo.17083860.

### Monensin Quantification

Monensin extractions were quantified using a vanillin assay. Dry extracts were reconstituted with methanol, mixed with vanillin straining solution (1:9 vanillin:MeOH sample/standard solution) and incubated 30 minutes at 55 °C and 180 rpm. Monensin reacts with the vanillin stain forming pink-coloured compounds that have an absorption maximum at 518-520 nm. The absorption spectra were measured for exactly 20 minutes in a SPECTROstar Nano microplate reader. Lambert-Beer’s law was used to determine the concentration of monensin inside the extracts using an external calibration curve (linearity in the range 0-200 mg/L, standards of monensin (Merck) were prepared in methanol. In general, dilutions were necessary for BPM extractions but not for the SM16 cultures.

### Premonensin Quantification

Fermentations, extractions, and reconstitutions were performed as described for monensin, with reconstitution volumes of 5 mL for SM16 cultures and 10 mL for BPM. A volume of 20 µL of methanol extracts was applied to TLC plates and developed using ethyl acetate:cyclohexane (6:4). Plates were stained with a vanillin solution (15 g vanillin, 2.5 mL concentrated H₂SO₄, 250 mL ethanol). Each plate contained four premonensin calibration standards as well as all media conditions from a single clone. Images were acquired using an Intas camera (black and white) with fixed distance and capture settings.

Densitometric analyses were carried out using LabImage 1D L340 (Intas Science Imaging GmbH). Background noise was removed using the “Minimum to Minimum” method, in which the background is estimated by connecting the minima of the pixel intensity profile. TLC plates used and the total titers per biological clone can be seen in Figure S21, and Table S7, respectively of the Supplementary Information A. All preprocessing steps in the Supplementary Information B (densitometric processing steps).

## Author Contributions

M.B. and F.S. performed sequence alignment and designed the mutations. S.K. carried out mutagenesis, including oligonucleotide design. J.H. conducted experiments and data analysis, including testing the effects of mutagenesis on premonensin and derivative titers, AlphaFold analyses, and qNMR. S.C.B. and F.S. designed the medium optimization and medium–mutation combination experiments for premonensin and monensin strains, which were performed by S.C.B. The manuscript was written by F.S, S.C.B., and J.H.

## Acknowledgments

We thank Max Küster for his contribution to the qNMR analyses of premonensin titers. We are grateful to Frank Peeters for his assistance with WT and BP monensin cloneś selection, optimization of monensin quantification, and medium effect experiments. We also thank Duc Thanh Dinh for his support in the oil effect and combination of medium and mutation experiments. S. C. B., J. H. and F. S. acknowledge funding from the German Research Foundation (DFG) through Research Training Group 2341 “Microbial Substrate Conversion (MiCon)”. S.C.B. is a fellow of the International Max Planck Research School for Living Matter: from molecules to dynamics (IMPRS-LM).

