## Supplementary Information for "Overcoming Biosynthetic Limitations to Enhance Bacterial Polyketide Production"

#### Introduction

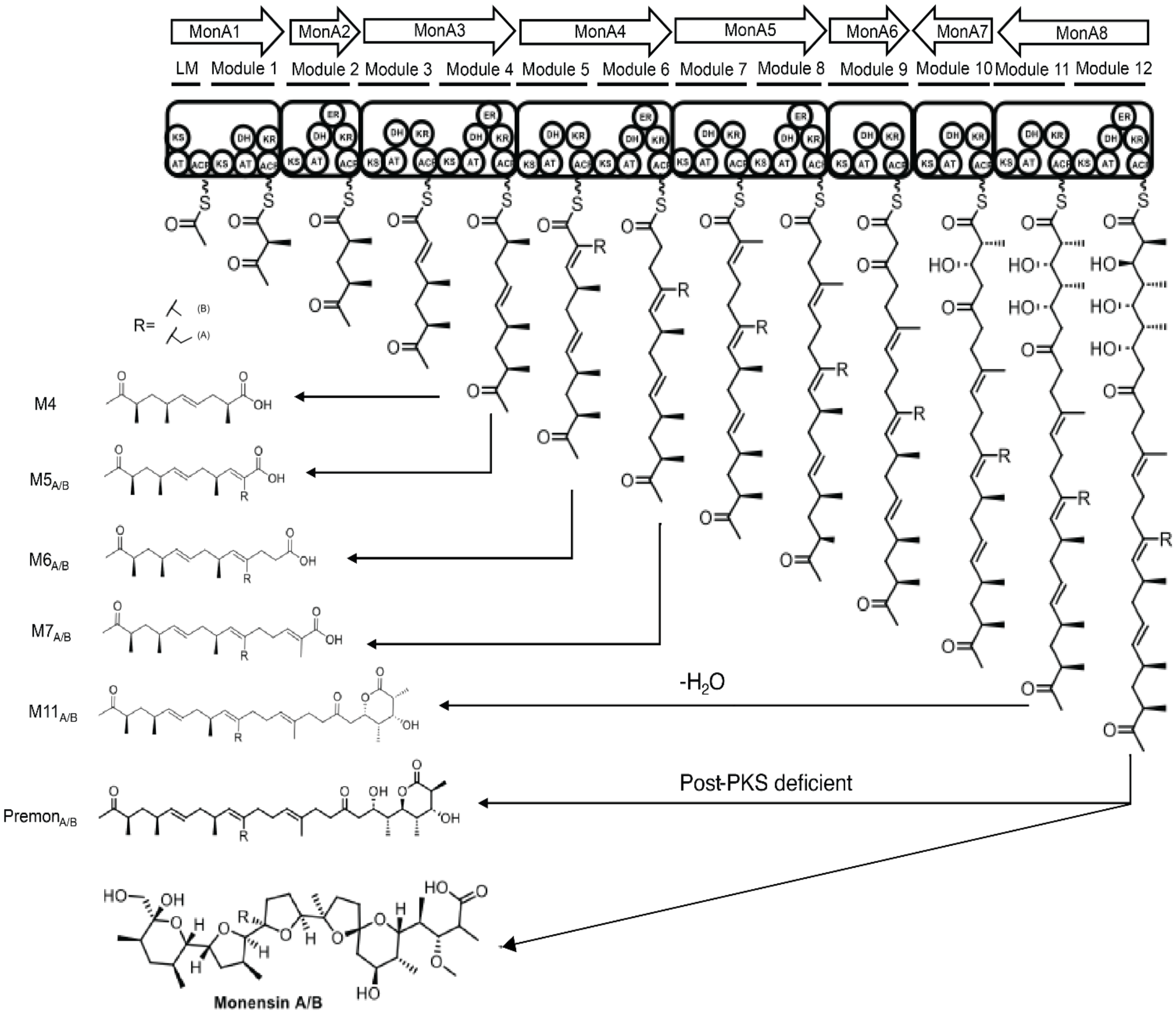


*Figure S1: Monensin is synthesized by a type I modular polyketide synthase (PKS), encoded by a biosynthetic gene cluster (BGC) comprising eight large proteins (MonAI–MonAVIII) that together harbor 12 modules. Each module contains the three core domains: ketosynthase (KS), acyltransferase (AT), and acyl carrier protein (ACP), and may additionally carry ketoreductase (KR), dehydratase (DH), and enoylreductase (ER) domains. This modular PKS machinery assembles a linear polyketide backbone, which is subsequently modified by a series of post-PKS tailoring enzymes to generate the final product, monensin. In Streptomyces cinnamonensis A495, deficiencies in certain post-PKS enzymes result in the accumulation of premonensin and shunt products (M4–M11). Due to the substrate promiscuity of the AT5 domain, both malonyl-CoA and methylmalonyl-CoA can be incorporated, giving rise to premonensin A and premonensin B.*

### PKS assembly line

#### Design and generation of single point mutants

##### A) Multiple Sequence Alignment of monensin KS domains

All monensin KS domain sequences (Accession ID: BGC0000100.5) were downloaded from the Minimum Information about a Biosynthetic Gene cluster (MIBiG) database(Zdouc et al. 2025). Multiple sequence alignment was performed using MUSCLE (Edgar 2004) with default parameters in UGENE v49.1.

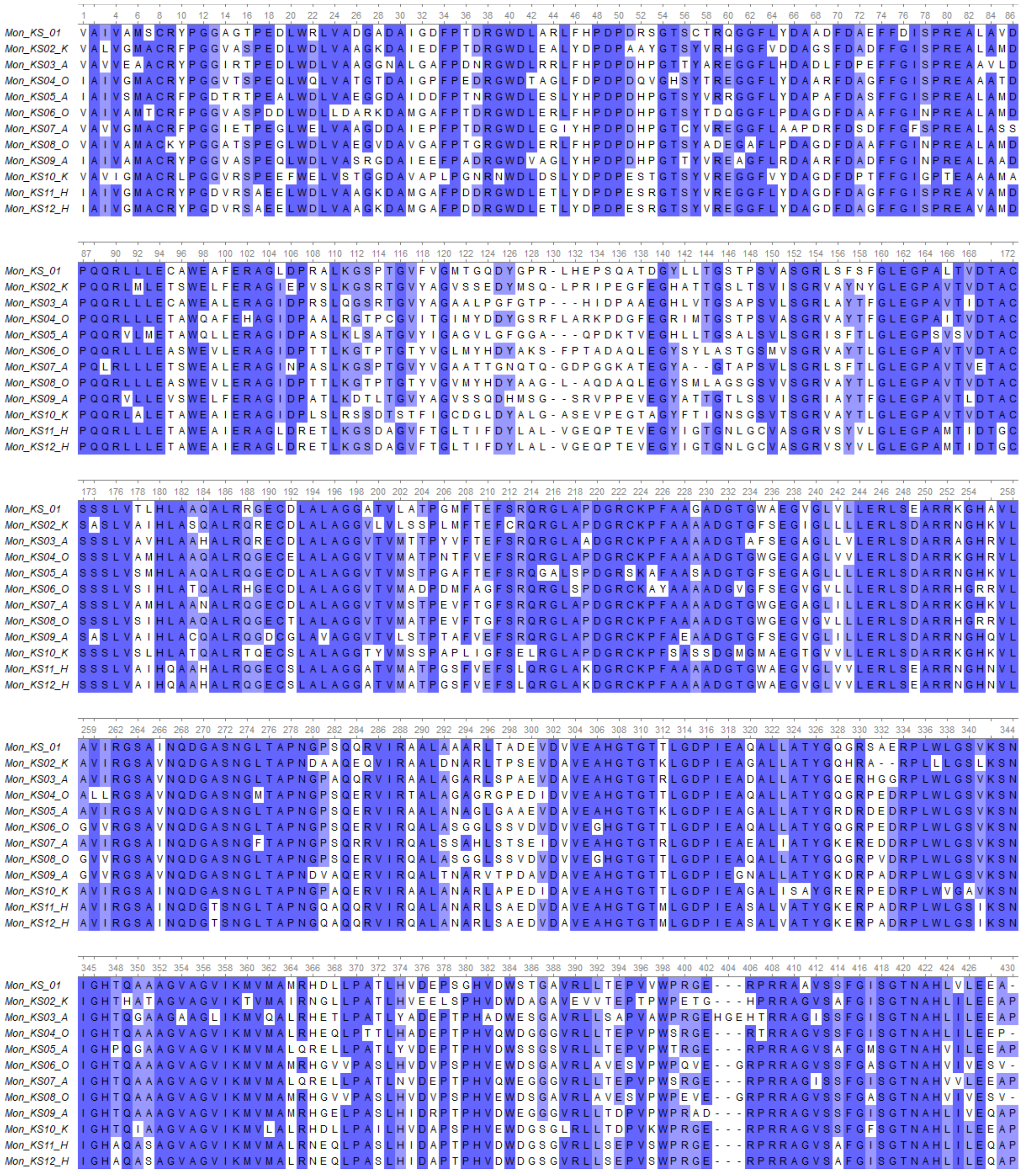

*Figure S2**:** Multiple sequence alignment of all ketosynthase domains from the monensin PKS. Residue conservation is highlighted at a threshold of ≥70%, and residues are color-coded based on percentage identity.*

#### B) Generation of S226A mutants

##### Plasmid construction

A *KS5* deletion construct was generated by replacing the *KS5* gene with a spectinomycin resistance (SptR) cassette. The construct was assembled by overlap extension PCR using fragments amplified with primer pairs 344–1978, 1977–345, and 723–724, and cloned into a HindIII- and EcoRI-digested pKC1139-based vector, carrying a temperature-sensitive replicon and apramycin resistance (AprR) cassette, via the SLIC-MIX method (Kushnir et al. 2012). To enable counter-selection, the *codA* gene from *E. coli* was inserted into the vector(Bierman et al. 1992). An identical procedure was used to generate plasmids carrying the S226A allele, using mutagenic primers 1963 and 1964.

##### Conjugation and selection

The resulting plasmids were first introduced into E. coli ET12567/pUZ8002 and then transferred into Streptomyces cinnamonensis A495 by conjugation. Selection of recombinants relied on the temperature-sensitive replicon.

In the first step, exconjugants were cultured at the non-permissive temperature (39 °C), where the plasmid cannot be replicated independently. Under these conditions, survival required a single homologous recombination event, integrating the plasmid into the chromosome. $Apr^R$ $Spt^R$ exconjugants were isolated at this stage.

In the second step, cultures were shifted back to the permissive temperature (30 °C) to allow a second homologous recombination event. This resolved the plasmid from the chromosome. Counter-selection with 5-fluorocytosine enriched for double-crossover recombinants, yielding $AprSSpt^R$ colonies that served as recipients for subsequent conjugations.

Plasmids carrying the S226A allele were then introduced into the recipient by conjugation, and $Apr^s$ $Spt^s$ variants were selected as described. As a control, the wild-type KS5 gene (amplified with primers 344–345) was cloned and reintroduced into the $AprSSpt^R$ recipient strain. After selection, suitable S colonies were identified by colony PCR and sequencing (oligos 1981-1982). Oligonucleotide sequences are shown in Table S1.

*Table S 1: Oligonucleotide used in this work*

| Oligo N | Oligo name | Oligo sequence |
|---|---|---|
| 344 | 344_KS5mut-1_for | atgacatgattacgaattctttgccgcggccgatgtcagtgatccggacgc |
| 345 | 345_KS5mut-2_rev | acggccagtgccaagctgtggtcgcggcgcagggtgggcacgaccgtgg |
| 723 | 723_res-marker_for | tgcagctcacggtaactgatgccgtatttgcagtaccagcg |
| 724 | 724_res-marker_rev | tggagctgcttcgaagttcctatactttctagagaataggaacttcgg |
| 1963 | 1963_KS5_AASA237-AAAA_for | acggccgctcgaaggctttcgcggccgcggccgacggcaccggtttctcggagggcgcgg |
| 1964 | 1964_KS5_AASA237-AAAA_rev | ccctccgagaaaccggtgccgtcggccgcggccgcgaaagccttcgagcggccgtccgg |
| 1977 | 1977_KS5-Swa_for | tggccatggacccgcagcagcgggtgctcatttaaatcaagatggtgatggcgctgcagcgcg |
| 1978 | 1978_KS5-Swa_rev | ttcgcgctgcagcgccatcaccatcttgatttaaatgagcacccgctgctgcgggtccatggcc |
| 1981 | 1981_KS5-seq1_rev | ttcctcgccccggccggagacgacccacgggacgactgcg |
| 1982 | 1982_KS5-seq2_for | aagcgggtcacggtggacctcggccaggcccgccggcgg |

*Table S 2: List of plasmids used for conjugation*

| N of Plasmid | Name | Source |
|---|---|---|
| 0079 | pKC1139 | (Breaud et al. 2022) |
| 1316 | pKC1139_codA#63 | This work |
| 1483 | pKC1139_codA#KS5-26_AASA237AAAA | This work |
| 1465 | pKC1139_codA#KS5-Swa | This work |
| 1437 | pKC1139_codA#KS5_sptR | This work |
| 1438 | pKC1139_codA#KS5wt | This work |

#### C) HPLC-MS Analysis

##### Spectra processing using Mzmine

All LC-MS *.d* files were converted to *.mzML* format using **MSConvert v3.0** (Chambers et al. 2012) and corrected using **Mzxml-Precursor-Corrector** (Breaud et al. 2022).

Followingly, all corrected data were imported into MZmine 3, and the batch mode module was used to process the data. The batch mode settings are as follows:

1) Mass detection for MS1 (noise level: 1000)
2) Mass detection for MS2 (noise level: 80)
3) Targeted feature detection (intensity tolerance: 20%, m/z tolerance: 10 ppm, retention time tolerance: 1 min).
4) Join aligner (m/z tolerance: 10 ppm, weight for m/z: 2, retention time tolerance: 1.5 min, weight for RT: 1
5) Group MS2 scans with features (MS1 to MS2 precursor tolerance: 20 ppm, retention time filter: use tolerance 0.2 min, minimum relative feature height: 25%, minimum required signal: 1)

##### MS1 and MS2 data of PreA/B, DH40PreA/B and shunt products

*Table S 3: List of characteristic adducts of PreA/B, DH4$^{0}$PreA/B and shunt products*

| | **meas. *m/z*** | **ion formula** | **calc. *m/z*** | **err [ppm]** |
|---|---|---|---|---|
| **PreB** | 583.3962 | $C_{34}H_{56}NaO_6$ | 583.3969 | 1.2 |
| | 599.3707 | $C_{34}H_{56}KO_6$ | 599.3708 | 0.2 |
| **PreA** | 597.4118 | $C_{35}H_{58}NaO_6$ | 597.4126 | 1.4 |
| | 613.385 | $C_{35}H_{58}KO_6$ | 613.3865 | 0.4 |
| **DH4$^{0}$PreB** | 599.3916 | $C_{34}H_{56}NaO_7$ | 599.3918 | 0.3 |
| **DH4$^{0}$PreA** | 613.4068 | $C_{35}H_{58}NaO_7$ | 613.4075 | 1.1 |
| **M4** | 249.1454 | $C_{13}H_{22}NaO_3$ | 249.1461 | 2.8 |
| **M5B** | 289.1794 | $C_{16}H_{26}NaO_3$ | 289.1774 | -6.9 |
| **M5A** | 303.1926 | $C_{17}H_{28}NaO_3$ | 303.1931 | 1.6 |

| | | | | |
|---|---|---|---|---|
| **M6B** | 317.2084 | $C_{18}H_{30}NaO_3$ | 317.2087 | 0.9 |
| **M6A** | 331.2242 | $C_{19}H_{32}NaO_3$ | 331.2244 | 0.6 |
| **M7A** | 349.2672 | $C_{22}H_{37}O_3$ | 349.2737 | 18.6 |
| | 371.2552 | $C_{22}H_{36}NaO_3$ | 371.2557 | 1.3 |
| **M11B** | 525.3541 | $C_{31}H_{50}NaO_5$ | 525.355 | 1.7 |
| **M11A** | 517.3881 | $C_{32}H_{53}O_5$ | 517.3888 | 1.4 |
| | 539.3701 | $C_{32}H_{52}NaO_5$ | 539.3707 | 1.1 |
| | 555.3445 | $C_{32}H_{52}KO_5$ | 555.3462 | 3.1 |

### PreA/B

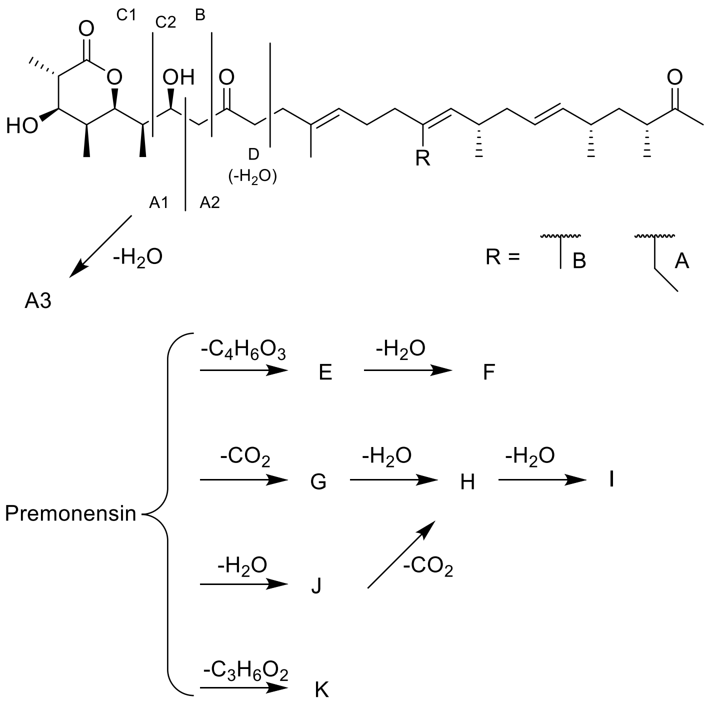


*Figure S3: Premonensin A/B structures and fragmentation patterns*

*Table S 4: Mass spectrometric data of the fragment ions of PreA/B.*

| Fragment | meas. *m/z* | ion formula | *m/z* | err [ppm] |
|---|---|---|---|---|
| | | **PreB** | | |
| **C1** | 193.0845 | $C_9H_{14}NaO_3$ | 193.0835 | -5.2 |
| **A3** | 205.0832 | $C_{10}H_{14}NaO_3$ | 205.0835 | 1.5 |
| **A1** | 223.0943 | $C_{10}H_{16}NaO_4$ | 223.0941 | -0.9 |
| **B** | 237.1087 | $C_{11}H_{18}NaO_4$ | 237.1097 | 4.2 |
| **D** | 263.1215 | $C_{13}H_{20}NaO_4$ | 263.1254 | 14.8 |
| **A2** | 383.2908 | $C_{24}H_{40}NaO_2$ | 383.2921 | 3.4 |
| **C2** | 411.2850 | $C_{25}H_{40}NaO_3$ | 411.287 | 4.9 |
| **F** | 463.3519 | $C_{30}H_{48}NaO_2$ | 463.3547 | 6.0 |
| **E** | 481.3650 | $C_{30}H_{50}NaO_3$ | 481.3652 | 0.4 |

| | | | | |
|---|---|---|---|---|
| **I** | 503.3853 | $C_{33}H_{52}NaO_2$ | 503.386 | 1.4 |
| **K** | 509.3513 | $C_{31}H_{50}NaO_4$ | 509.3601 | 17.3 |
| **J** | 521.3954 | $C_{33}H_{54}NaO_3$ | 521.3965 | 2.1 |
| **G** | 539.4025 | $C_{33}H_{56}NaO_4$ | 539.4071 | 8.5 |
| **H** | 565.3839 | $C_{34}H_{54}NaO_5$ | 565.3863 | 4.2 |
| | | **PreA** | | |
| **C1** | 193.0839 | $C_9H_{14}NaO_3$ | 193.0835 | -2.1 |
| **A3** | 205.0837 | $C_{10}H_{14}NaO_3$ | 205.0835 | -1.0 |
| **A1** | 223.0944 | $C_{10}H_{16}NaO_4$ | 223.0941 | -1.3 |
| **B** | 237.1112 | $C_{11}H_{18}NaO_4$ | 237.1097 | -6.3 |
| **C2** | 263.1261 | $C_{13}H_{20}NaO_4$ | 263.1254 | -2.7 |
| **A2** | 397.3071 | $C_{25}H_{42}NaO_2$ | 397.3077 | 1.5 |
| **D** | 425.3012 | $C_{26}H_{42}NaO_3$ | 425.3026 | 3.3 |
| **F** | 477.3704 | $C_{31}H_{50}NaO_2$ | 477.3703 | -0.2 |
| **E** | 495.3827 | $C_{31}H_{52}NaO_3$ | 495.3809 | -3.6 |
| **I** | 517.4014 | $C_{34}H_{54}NaO_2$ | 517.4016 | 0.4 |
| **K** | 523.3772 | $C_{32}H_{52}NaO_4$ | 523.3758 | -2.7 |
| **J** | 535.4060 | $C_{34}H_{56}NaO_3$ | 535.4122 | 11.6 |
| **G** | 553.4148 | $C_{34}H_{58}NaO_4$ | 553.4227 | 14.3 |
| **H** | 579.3977 | $C_{35}H_{56}NaO_5$ | 579.402 | 7.4 |

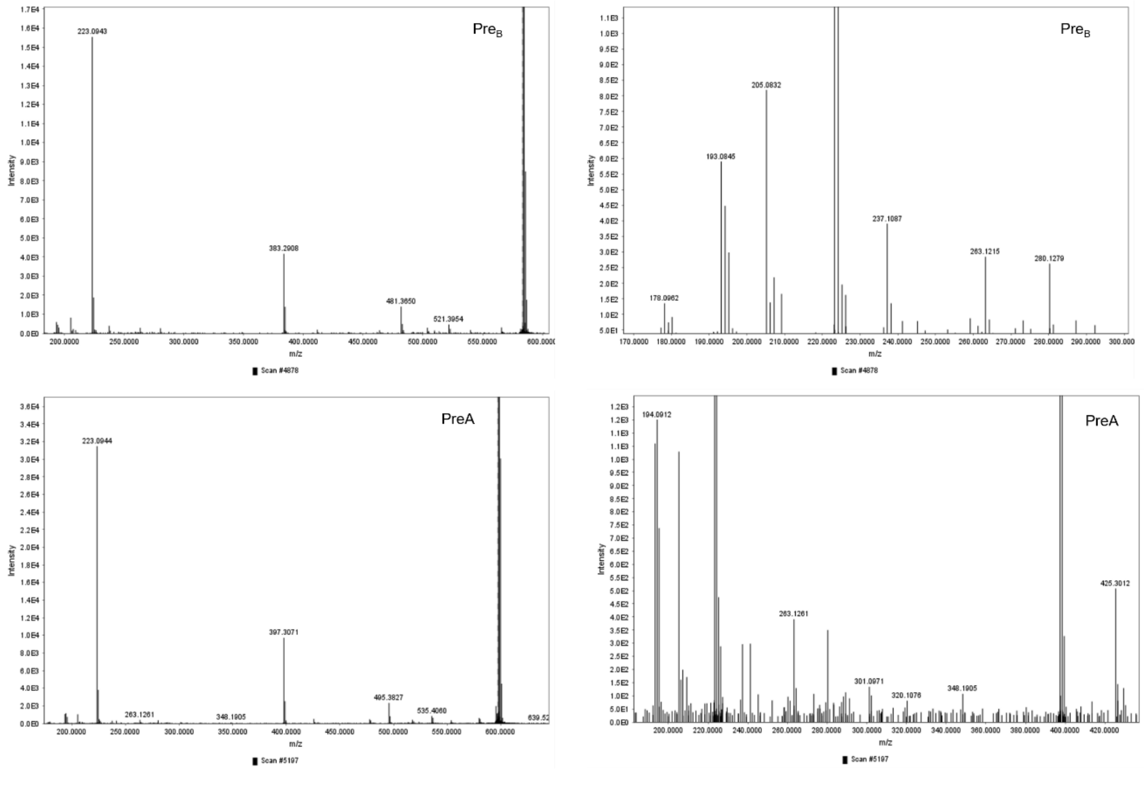


*Figure S4: MS² spectra of PreA/B.*

### DH40PreA/B

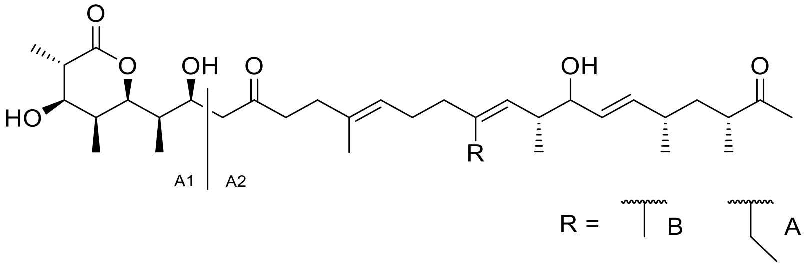


*Figure S5: Fragmentation pattern of DH4$^0$PreA/B*

*Table S 5: Mass spectrometric data of the fragment ions of DH4$^0$PreA/B.*

| Fragment | meas. m/z | ion formula | m/z | err [ppm] |
|---|---|---|---|---|
| | | **DH4$^0$PreB** | | |
| **A1** | 223.0956 | $C_{10}H_{16}NaO_4$ | 223.0941 | 0.3 |
| **A2** | 399.2855 | $C_{24}H_{40}NaO_3$ | 399.2870 | 1.3 |
| | | **DH4$^0$PreA** | | |
| **A1** | 223.0931 | $C_{10}H_{16}NaO_4$ | 223.0941 | 1.3 |
| **A2** | 413.3023 | $C_{25}H_{42}NaO_3$ | 413.3026 | 2.3 |

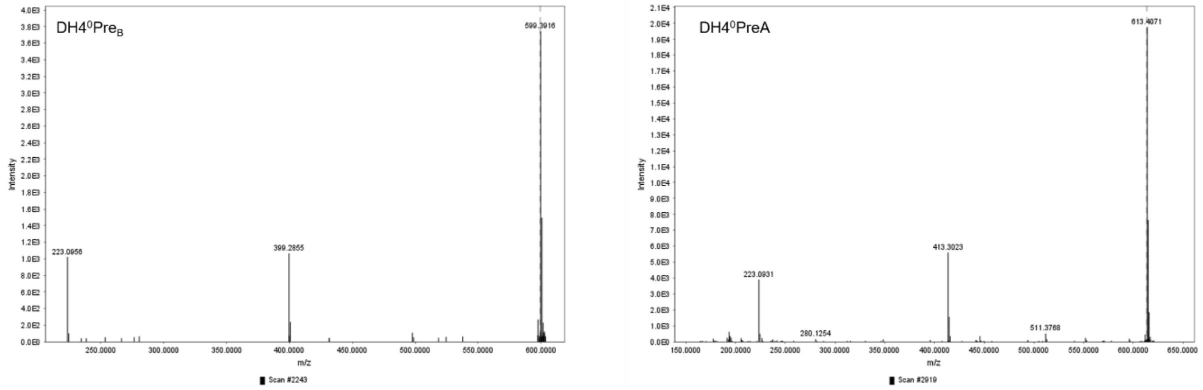


*Figure S6: MS$^2$ spectra of DH4$^0$PreA/B.*

### Shunt Products

Only the MS² fragmentation patterns of M11A/B are shown, as the other compounds did not fragment under the applied conditions.

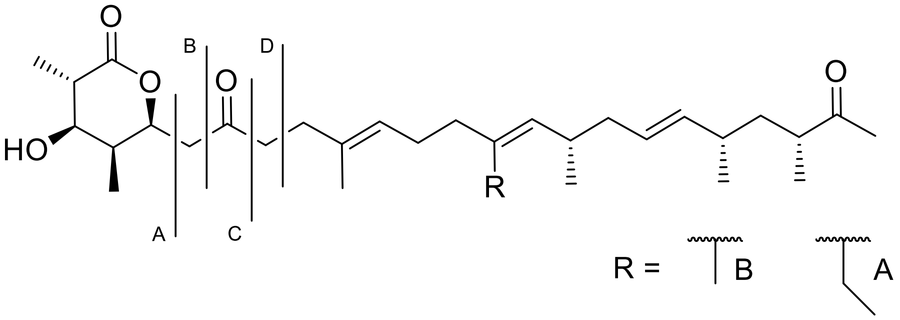


*Figure S7: Fragmentation pattern of M11A/B*

*Table S 6: Mass spectrometric data of the fragment ions of M11A/B*

| Fragment | Meas. m/z | ion formula | m/z | err [ppm] |
|---|---|---|---|---|
| | | **M11B** | | |
| **A** | 165.052 | $C_7H_{10}NaO_3$ | 165.0522 | 1.2 |
| **B** | 179.0669 | $C_8H_{12}NaO_3$ | 179.0679 | 5.6 |
| **C** | 207.0624 | $C_9H_{12}NaO_4$ | 207.0628 | 1.9 |
| **D** | 222.0841 | $C_{10}H_{15}NaO_4$ | 222.0863 | 9.9 |
| | | **M11A** | | |
| **A** | 165.0509 | $C_7H_{10}NaO_3$ | 165.0522 | 7.9 |
| **B** | 179.0684 | $C_8H_{12}NaO_3$ | 179.0679 | -2.8 |
| **C** | 207.0627 | $C_9H_{12}NaO_4$ | 207.0628 | 0.5 |
| **D** | 222.0851 | $C_{10}H_{15}NaO_4$ | 222.0863 | 5.4 |

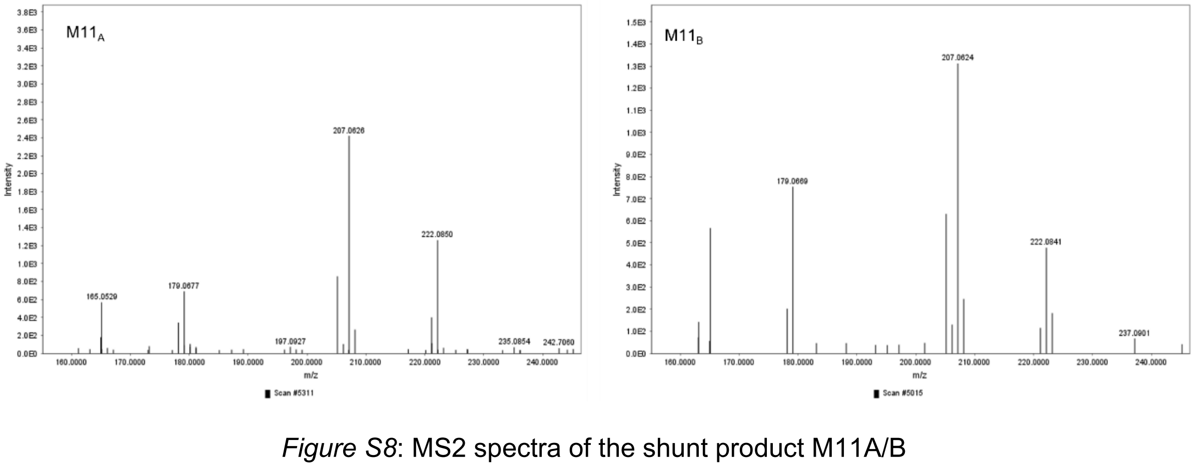


*Figure S8*: MS2 spectra of the shunt product M11A/B

#### D) AlphaFold3 structure prediction

Models for structure visualization were generated by AlphaFold3 Beta server. The following amino acid sequences were uploaded into the server (entity type: protein; copies: 2)

**ACP4DD-DDKS5-WT-Model0 sequence:**

>LAGLPHPERRRLLLDLVRGNVAGVLGHSDHDAVRPDTSFKELGFDSLTAVELRNR
LAAATGLKLPAALVFDYPESATLVDHLLERLPVLNDLGRIESSLDALALDADARSRVT
RRLNTLLSKLNGAATAGSPADVTDLDALDALDDVSDDEMFEFIDRELEAKLRQYLK
RVTVDLGQARRRLREVEERIAIVSMACRFPGDTRTPEALWDLVAEGGDAIDDFPTN
RGWDLESLYHPDPDHPGTSYVRRGGFLYDAPAFDASFFGISPREALAMDPQQRVL
METAWQLLERAGIDPASLKLSATGVYIGAGVLGFGGAQPDKTVEGHLLTGSALSVL
SGRISFTLGLEGPSVSVDTACSSSLVSMHLAAQALRQGECDLALAGGVTVMSTPG
AFTEFSRQGALSPDGRSKAFAASADGTGFSEGAGLLLLERLSDARRNGHKVLAVIR
GSAVNQDGASNGLTAPNGPSQERVIRAALANAGLGAAEVDAVEAHGTGTKLGDPI
EAGALLATYGRDRDEDRPLWLGSVKSNIGHPQGAAGVAGVIKMVMALQRELLPAT
LYVDEPTPHVDWSSGSVRLLTEPVPWTRGERPRRAGVSAFGMSGTNAHVILEEAP

**ACP4DD_DDKS5_S226A-Model0 sequence:**

>LAGLPHPERRRLLLDLVRGNVAGVLGHSDHDAVRPDTSFKELGFDSLTAVELRNR
LAAATGLKLPAALVFDYPESATLVDHLLERLPVLNDLGRIESSLDALALDADARSRVT
RRLNTLLSKLNGAATAGSPADVTDLDALDALDDVSDDEMFEFIDRELEAKLRQYLK
RVTVDLGQARRRLREVEERIAIVSMACRFPGDTRTPEALWDLVAEGGDAIDDFPTN
RGWDLESLYHPDPDHPGTSYVRRGGFLYDAPAFDASFFGISPREALAMDPQQRVL
METAWQLLERAGIDPASLKLSATGVYIGAGVLGFGGAQPDKTVEGHLLTGSALSVL
SGRISFTLGLEGPSVSVDTACSSSLVSMHLAAQALRQGECDLALAGGVTVMSTPG
AFTEFSRQGALSPDGRSKAFAAAADGTGFSEGAGLLLLERLSDARRNGHKVLAVIR
GSAVNQDGASNGLTAPNGPSQERVIRAALANAGLGAAEVDAVEAHGTGTKLGDPI
EAGALLATYGRDRDEDRPLWLGSVKSNIGHPQGAAGVAGVIKMVMALQRELLPAT
LYVDEPTPHVDWSSGSVRLLTEPVPWTRGERPRRAGVSAFGMSGTNAHVILEEAP

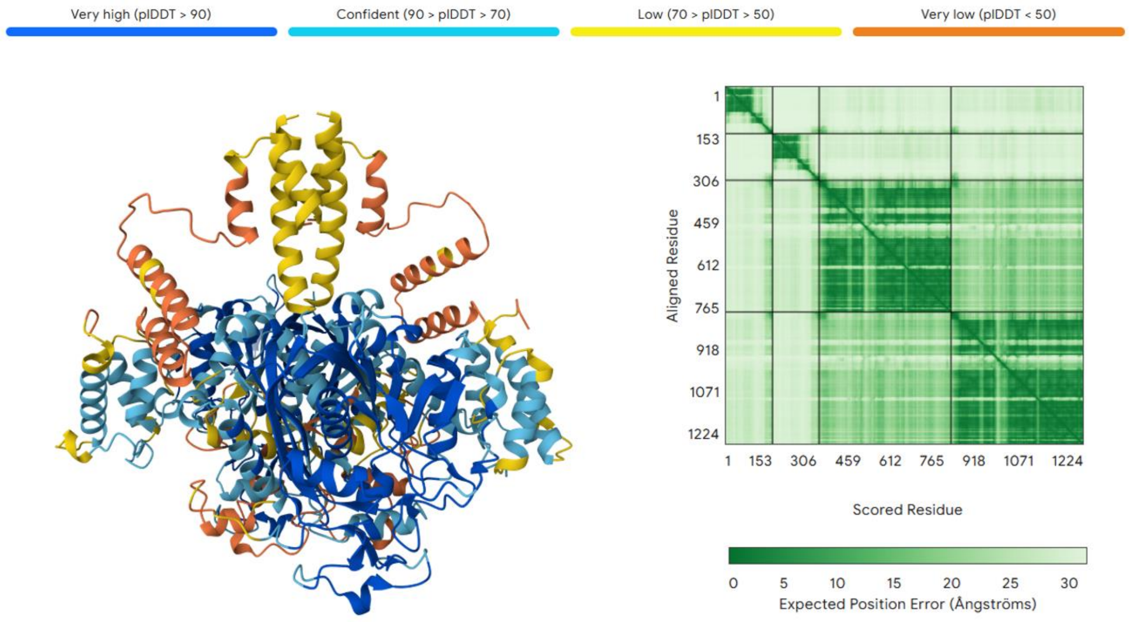


*Figure S9: Alpha fold 3 beta model of WT-ACP4-$^{C}$docking domain4 and $^{N}$docking domain5-KS5 homodimers. The ribbon diagram (left) is colored by per-residue confidence scores (pLDDT), with blue indicating very high confidence (pLDDT > 90) and orange indicating very low confidence (pLDDT < 50). The predicted aligned error (PAE) heatmap (right) represents the expected positional error in Ångströms between residue pairs, with darker green indicating higher certainty. Overall prediction confidence is summarized by predicted TM-score (pTM = 0.54) and interface predicted TM-score (ipTM = 0.33).*

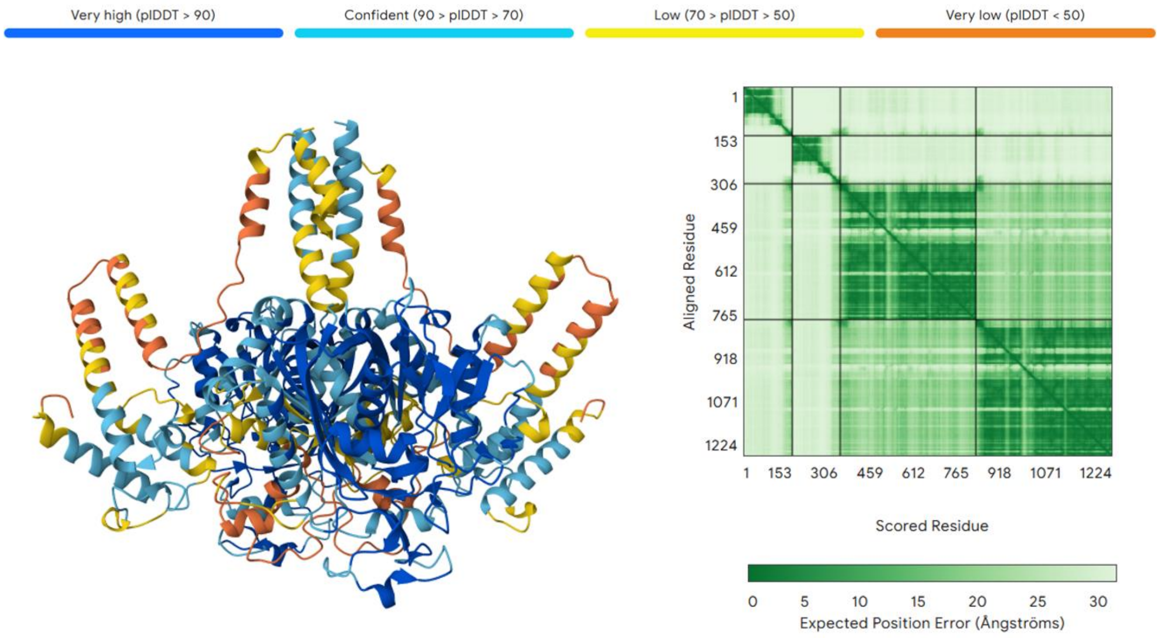


*Figure S10**:** Alpha fold 3 beta model of S226A-ACP4-$^{C}$docking domain4 and $^{N}$docking domain5-KS5 homodimers. The ribbon diagram (left) is colored by per-residue confidence scores (pLDDT), with blue indicating very high confidence (pLDDT > 90) and orange indicating very low confidence (pLDDT < 50). The predicted aligned error (PAE) heatmap (right) represents the expected positional error in Ångströms between residue pairs, with darker green indicating higher certainty. Overall prediction confidence is summarized by predicted TM-score (pTM = 0.53) and interface predicted TM-score (ipTM = 0.32).*

### Media Optimization in Monensin Production

#### E) Quality control and optimization of Monensin Quantification

The absorption spectra were measured using a SPECTROstar® Nano microplate reader with the following settings: endpoint, 30 second shake before measurement, 20 pulses, 520 nm

For each plate of measurements, twelve concentrations monensin standards (in duplicates) were measured in simultaneous. Standard solutions were prepared by duplicate, starting from a 1000 mg/L monensin in methanol stock solution as follows. 0, 10, 20, 25, 50, 75, 100, 125, 150, 175, 200 and 250 mg/L. Linearity was conserved in the 0-200 mg/L ($r^2$>0.93, in all cases). An overlap of these spectra is depicted on Figure S11. The absorption spectrum of a BPM medium blank is also added and it has a minimal absorption at 518 nm, whereas the absorption of dilution samples increases with increasing concentration of monensin, clearly indicating that monensin is responsible for absorption at 518 nm. Furthermore, when the BPM absorption level is considered the noise level on the right), absorption of different colonies normally exceeded the minimal S/N ratio of 3.

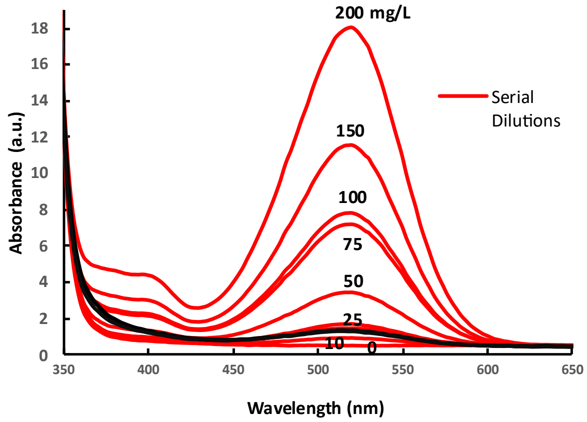


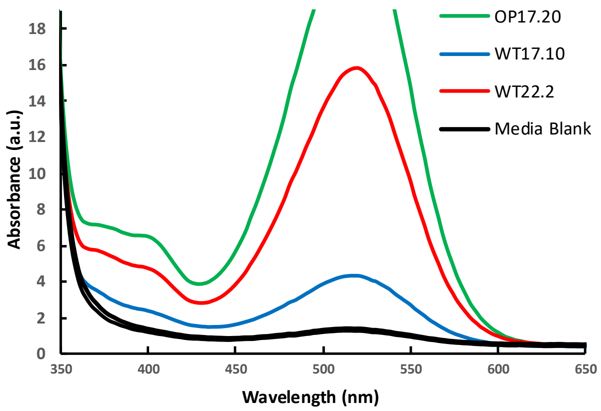


*Figure S11: (Left) Absorption spectra solutions (red) of different concentrations of monensin in MeOH as well as BPM media blanks (black). Media blank absorption at 518 nm corresponds to approx. 20 mg/L monensin in MeOH (Right): the same media blanks are plotted together with spectra of selected cultures. A good S/N ratio was observed in most cases.*

The time duration between incubation of the organic phase with the vanillin mixture and preparing the 96-well plate turned out to influence the slope of the calibration curve that was later obtained from the absorptions of the dilution row samples. A time window of 20 minutes was determined to be a good compromise between assay sensibility, robustness and good linearity.

Absorbances from the sample dilutions were used to obtain concentration according to the calibration curve. The use or not of dilutions was to make sure all samples could be interpolated from the calibration line. Values obtained by the curve were corrected according to the applied dilution and to the original fermentation medium volume and methanol reconstitution volume used.

**Monensin Cross validation Assays**

Accurate and efficient quantification of monensin is pivotal for both maintenance and optimization of media and growth conditions.

To identify the most suitable approach, we systematically evaluated multiple quantification methods, including direct High Pressure Liquid Chromatography-Mass Spectrometry (HPLC-MS or, LC-MS for short) of extracts, as well as indirect methods such as colorimetric quantification using vanillin staining, and antibiotic activity assays. Due to the different unit outputs of these methods, we employed median normalization for inter-method comparisons.

The inhibition zone method, although offering a labor-efficient approach with lower hands-on working time compared to other methods, is prone to false positive readouts from co-produced metabolites. Conversely, LC-MS analysis, notwithstanding its high sensitivity, is limited by incompatibility with extracts from fermentation media with oil additives. In contrast, the vanillin assay demonstrated reliability and reproducibility across diverse media.

Extensive cross-validation across multiple clones and media formulations confirmed the vanillin assay to be best suitable for routine monensin production assessment, owing to its quantitative nature and high-throughput compatibility. Although this approach is labour-intensive, its overall advantages make it the preferred choice for quantifying monensin production.

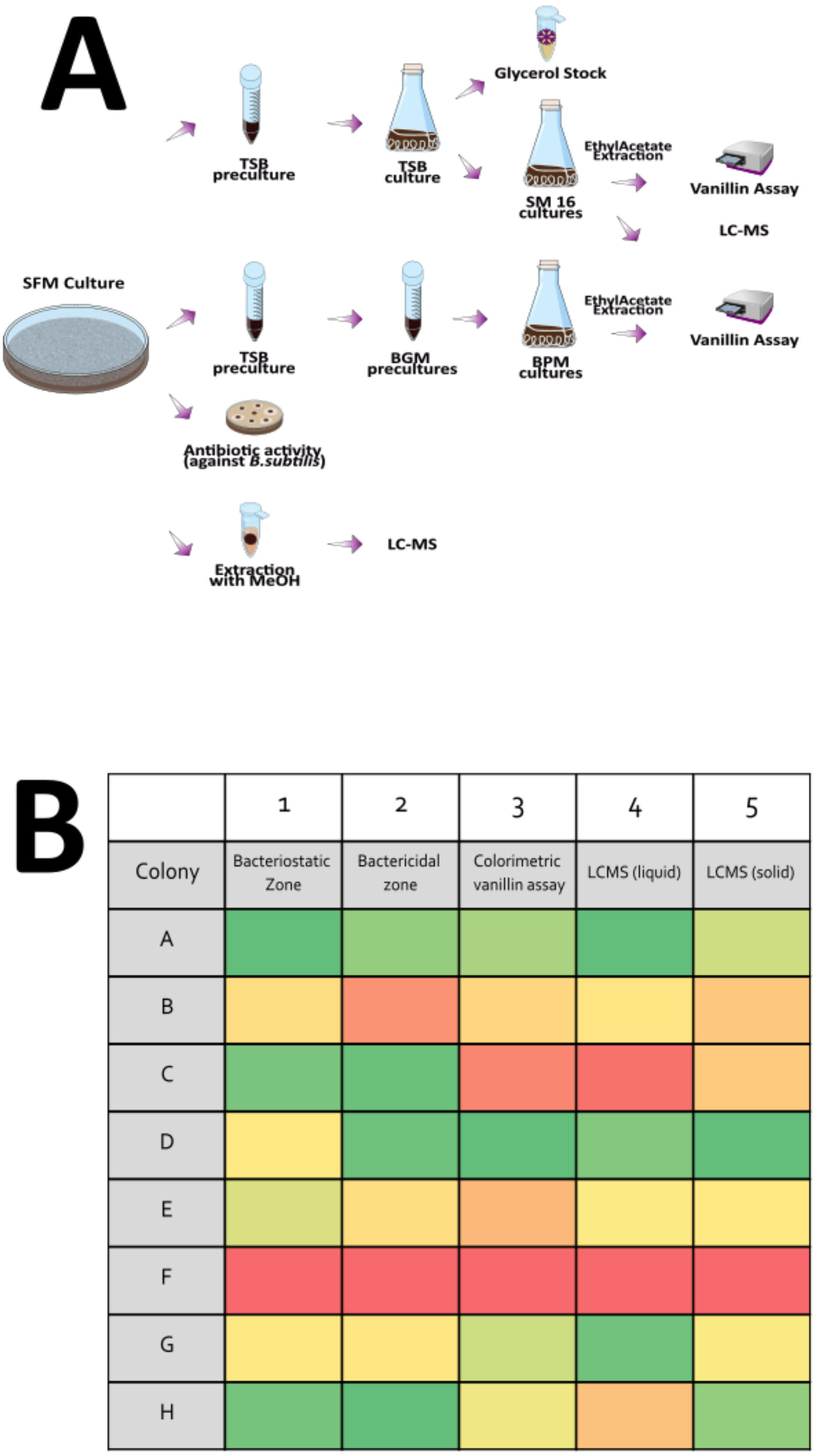


*Figure S12: Comparative analysis of monensin production across multiple clones of the Bulgarian Producer variant of Streptomyces cinnamonensis (BP). A. Experimental Workflow and Sample Preparation: Clones A-H were isolated from serial dilutions of spores of the BP stock. Each clone was subsequently cultivated on a separate SFM agar plate for 7 days. Standardized agar plugs served as inocula for TSB precultures and were used for inhibition assays using Bacillus subtilis. Monensin titers in SM16 and Bulgarian Production Medium (BPM) fermentations were quantified via vanillin assay. Methanolic extracts from both agar plugs and SM16 cultures were further analyzed by LCMS. B. Heatmap of normalized monensin production values data from 8 independent biological replicates were normalized and visualized as a heatmap.*

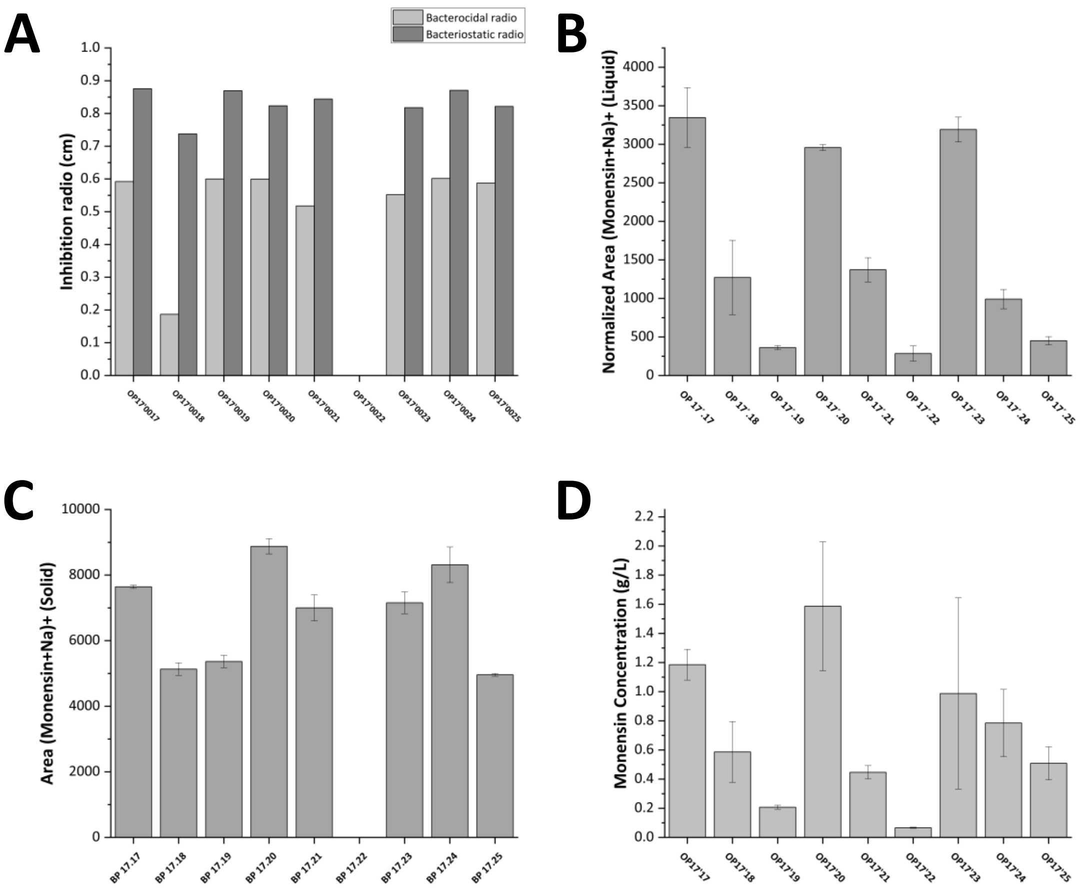


*Figure S13: Individual Measurements of each cross validation assay. Error bars correspond to the standard deviation. A) Inhibition radio of inner halo of measurements. B) Normalized area of monensin sodium adduct extracted ion chromatogram of liquid cultures (SM16). C) Normalized area of monensin sodium adduct extracted ion chromatogram of solid cultures (SFM). D) Monensin concentration according to the Vanillin Assay.*

#### Relative quantification of monensin in SFM agar plugs through mass spectrometry

Industrial strain BP clones were grown on SFM agar petri dishes for a week, a P1000 tip size plug, collected with the aid of sterilized forceps and toothpicks, transferred to 2 ml Eppendorf tubes, and frozen at -80°C for 20 min. Consecutively, 1 ml of LCMS-grade MeOH was added, vortexed for 15 min (2500 rpm pulse mode, RT) and centrifuged. 800 µl of the supernatant was isolated and frozen at -80°C for 30 min. Afterwards, the samples were centrifuged (12000 rpm, 30 min; 4 °C) to get particle-free solution for the HPLC measurements. Samples were diluted in an HPLC-vial: 10 µl of the previous supernatant in 990 µl of LCMS grade MeOH. Direct injections of the sample were measured in positive ion mode. The HPLC run lasted 3 min and it was isocratic 75% acetonitrile (0.1% FA): 25% water (0.1% FA), MN HPLC EC 4/2 Universal RP. Injection blanks were injected between samples to prevent carryover effects. LCMS data were processed in MZMine3 (peak) and the total area of the XIC's of monensin. For relative quantification, the sodium adduct area was used (RT: 0.4 min, m/z 693.4215).

**Relative quantification of Monensin in SM-16 cultures through direct injections.**

From each culture, an aliquot of 1 ml was transferred to a 1.5 ml Eppendorf tube and frozen at -80 °C. After the sample had thawed, it was centrifugated (5 min, 4°C, 5500 rpm), 800 µl of the supernatants were taken and transferred to new Eppendorf tubes and freeze-dried. The samples were reconstituted in 1 ml of LCMS-grade MeOH and then prepared for the HPLC. The samples were frozen to induce precipitation of low solubility molecules (monensin is very soluble in methanol). Particulates were eliminated by centrifugation (15 min, 4°C, 12700 rpm), each sample was diluted 1:100 with LCMS-grade MeOH inside a HPLC-vial. Samples were analyzed exactly the same as with the solid plugs. For quantification, the sodium adduct area was used (RT: 0.4 min, m/z 693.4184).

#### Inhibition zone analysis

500 µl of a B. subtilis stock solution was added to an autoclaved Erlenmeyer flask containing 150 ml of melted LB agar. Petri dishes were prepared with 15 ml culture mix, allowed to cool and solidify inside the fridge (4°C) for 40 min. With sterilized tweezers a 1 ml Eppendorf pipette tip size *S. cinnamonensis* plugs were taken from each week grown SFM culture and placed on top of the LB agar plates that contained B. subtilis. Five plugs were put on one plate and a record was made of their respective location. The prepared plates were incubated at 30°C for two days, and pictures of the plates were made using an iX imager of INTAS science imaging. A ruler was part of each of the photos for calibration of the size of the inhibition zones. Using ImageJ, the

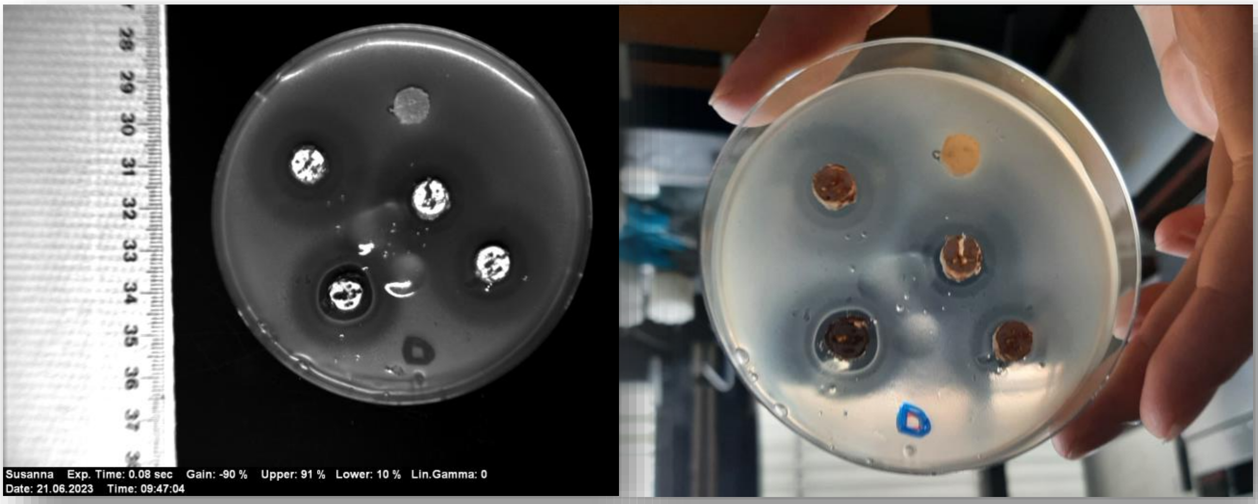


*Figure S14: Antibiotic activity assay. Five samples were tested per plate, each sample was tested in triplicate. (Left) picture by INTAS software. (Right):Real look of the inhibition test to the naked eye Two types of inhibition halos were observed. The ones used for the crossvalidation were the inner halos.*

zones of inhibition were measured. In steps of 45°, the length of the inhibition zone was measured giving eight values per zone. An average was taken of the eight measurements, and the standard deviation was calculated as an estimation of the assay standard deviation. (Exposure time 0.12 s, Gain: -80%, Upper contrast: 91%, Lower contrast: 5%, Linear gamma: 0).

#### F) Selection of best clones

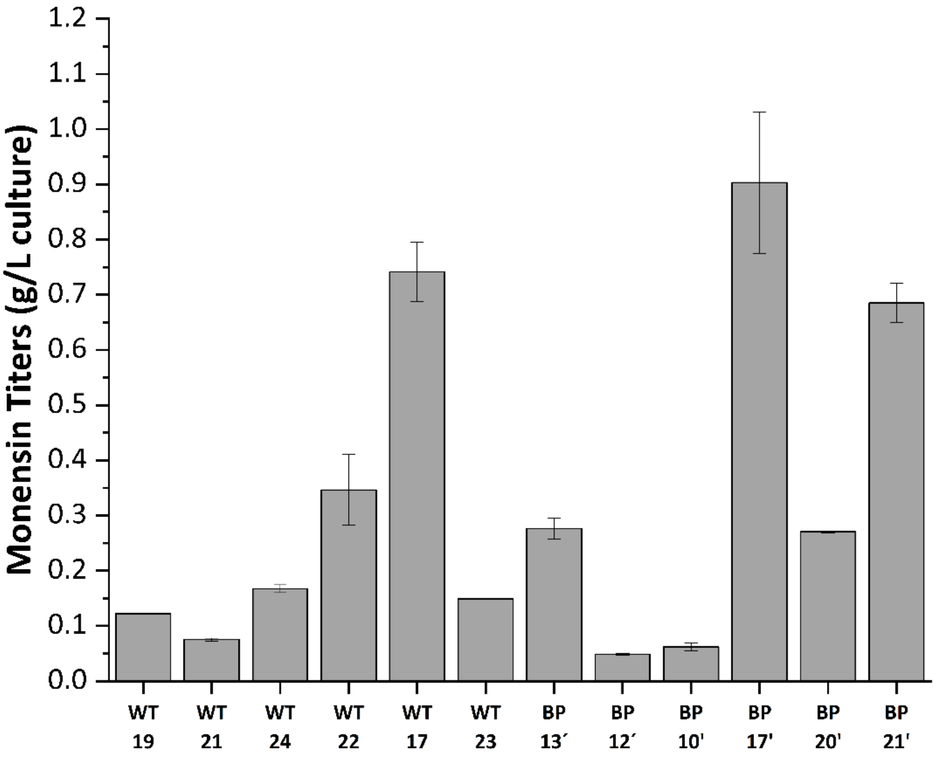


*Figure S15: First round of selection of clones. Bars show monensin production in Bulgarian Production Medium using Vanillin Assay. Values are average from extraction duplicates and error bars are standard deviation from each sample. Clones WT 22.2, WT 17.10 and BP 17.20 came from WT 22, WT 17 and BP 17, respectively.*

#### G) Cell Dry Weight (CDW) analysis:

Three aliquots of 1 ml of each culture were transferred to a tared microcentrifuge tube, spinned down and washed three times with distill water. The samples were freezedried and CDW was calculated.

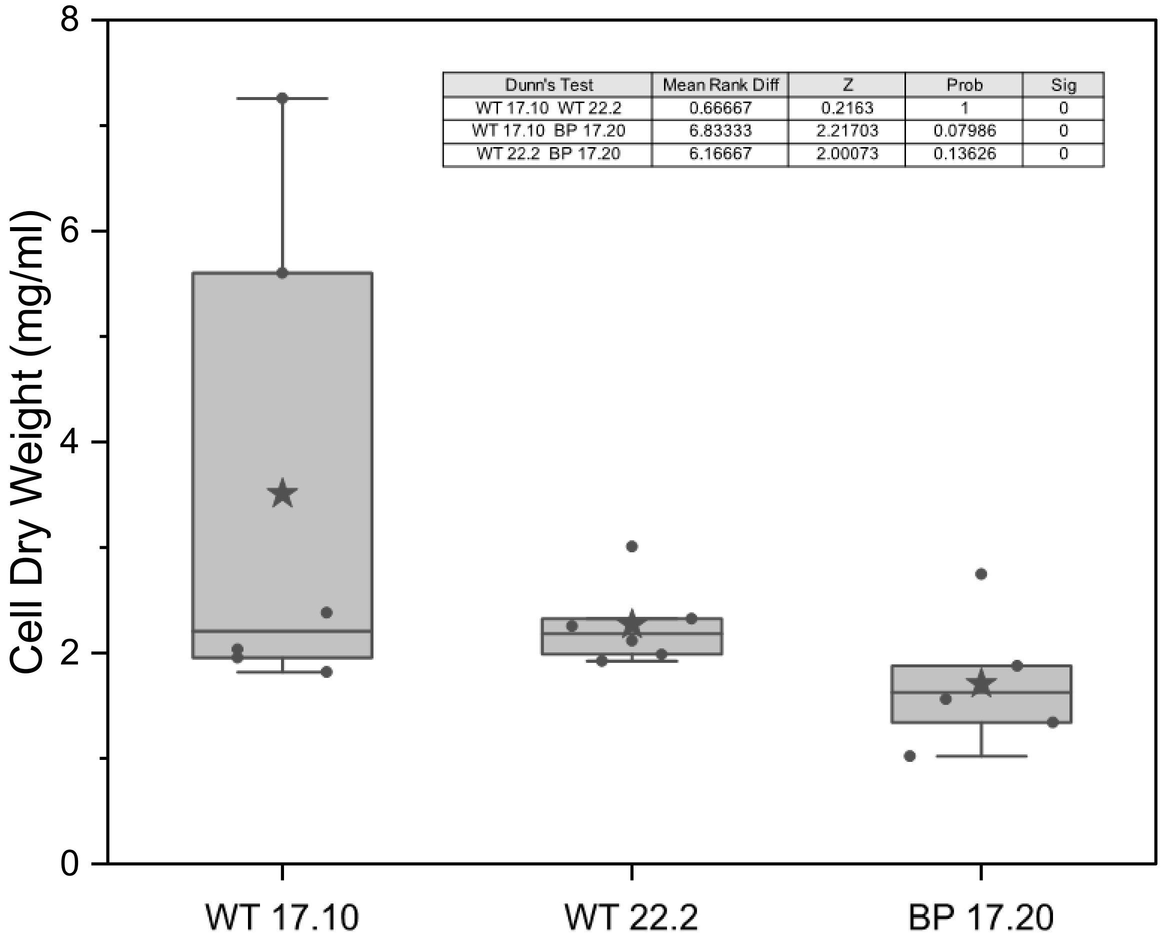


*Figure S16: Cell dry weight measurements of SM16 cultures. Triplicates were used to calculate each CDW. The average of each biological replicate is shown with black dots. The mean of each clone is marked with a black star, the median with a line. Non parametric tests were applied to the groups and results are shown in the table. There was no statistically significat difference between the CDW of the clones.*

#### H) Exploring Medium Variations for Enhanced Production

##### Effect of the Oil Composition

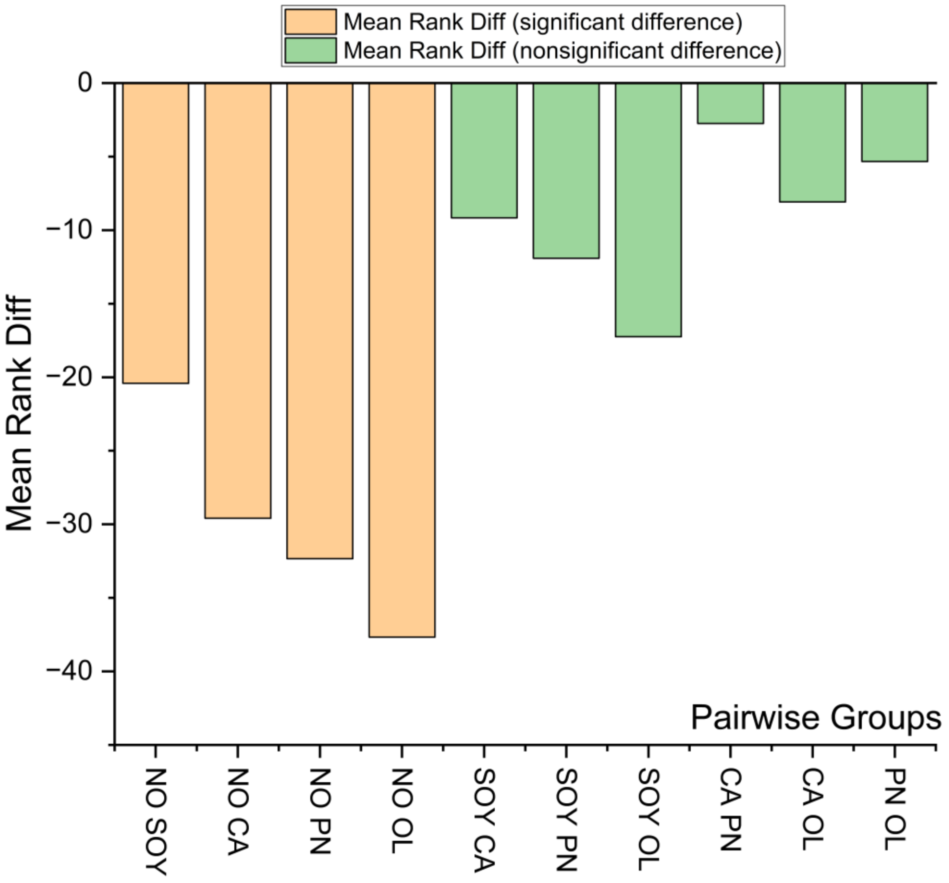


*Figure S17: Mean rank differences between different oil groups. Kruskal–Wallis with Dunn's post-hoc test was used, where raw values were converted into ranks. The Mean Rank Difference represents the difference between the average ranks of two groups. Here, No Oil (NO), Soybean oil (SOY), Canola oil (CA), Peanut oil (PN), and Olive oil (OL) were compared. Samples are paired, so, for example, 'NO SOY' represents the comparison between no oil and soybean oil. Negative values indicate that the first group (e.g., NO) has lower values than the second (e.g., SOY). Values closer to zero indicate more similar distributions, while larger positive or negative values reflect greater differences in raw values. There was a significant increase when using any oil compared to no oil, but no significant differences (Non parametric tests, Dunn´s, Origin 2024b) were found among the oils, although olive oil showed the strongest trend.*

#### Effect of the Additives

The additions were tested in both SM16 and BPM in a 5-day-cultivation, extracted and quantified with the standard operating procedure of the Vanillin Assay, with a monensin external calibration curve.

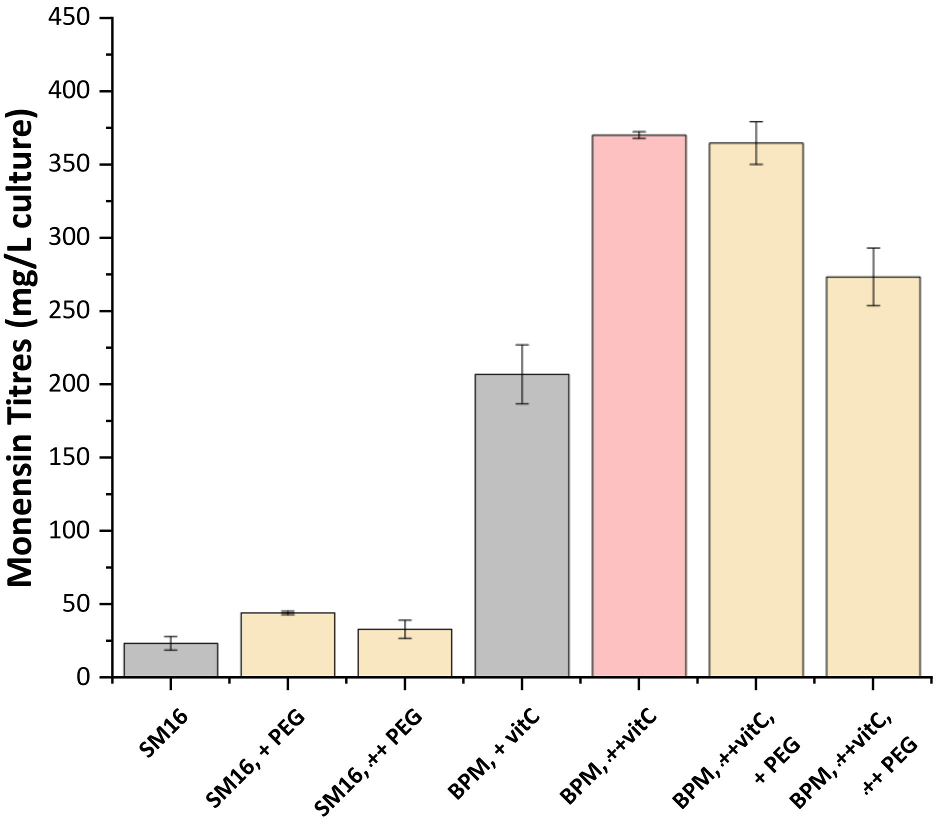


*Figure S18: Effect of PEG and high ascorbic acid in SM16 and BPM cultures. The values are the average from two biological replicates with two technical replicates each, the error bars come from the standard deviation of each. Plus (+) symbols refer to concentration tested (+PEG = 5 g/L, ++PEG= 10 g/L PEG, +vitC= 19 mg/L, ++vitC=110 mg/L).*

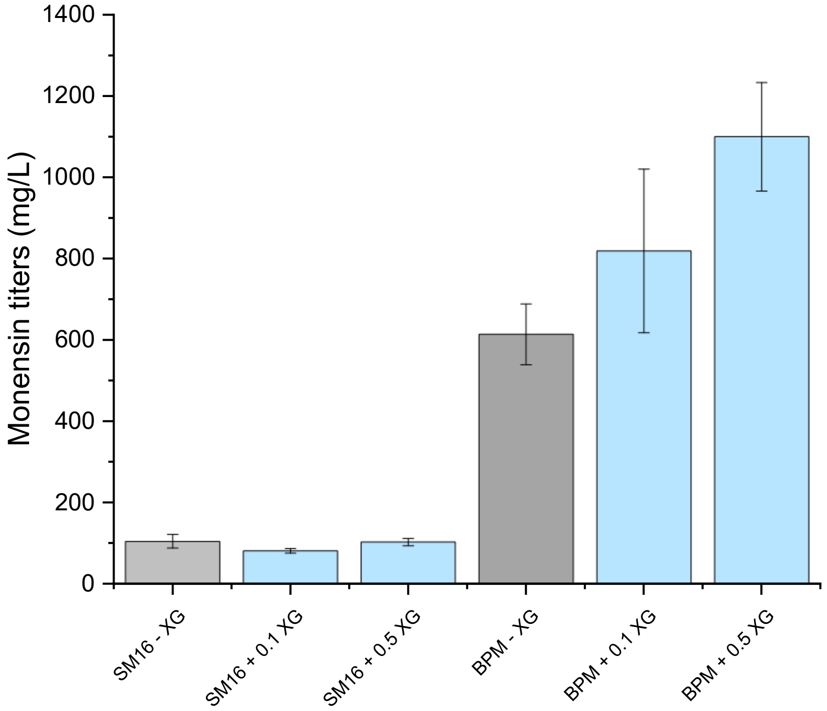


*Figure S19: Effect of Xanthan Gum (XG) in SM16 and BPM cultures. The values are the average from two biological replicates with two technical replicates each, the error bars come from the standard deviation of each. Substract sign (-) refers to no addition. Plus (+) symbols refer to concentration tested (+0.1 = 0.1 g /L, +0.5= 0.5 g/L XG).*

#### Beyond the norm: Testing the Medium Effect on premonensin

The vanillin assay, initially developed for monensin, was adapted for premonensin quantification, involving optimized temperatures and incubation times. The assay linearity was verified using purified premonensin, with a concentration range of 0-2000 mg/L, as confirmed by qNMR. However, matrix effects were observed in the use of crude fermentation extracts for spectrophotometric analysis, which were mitigated by thin-layer chromatography (TLC) analysis. TLC visualization of standards proved to be more reliable method than spectrophotometric measurements, as many compounds absorbing in the same range could be separated in the rf dimension.

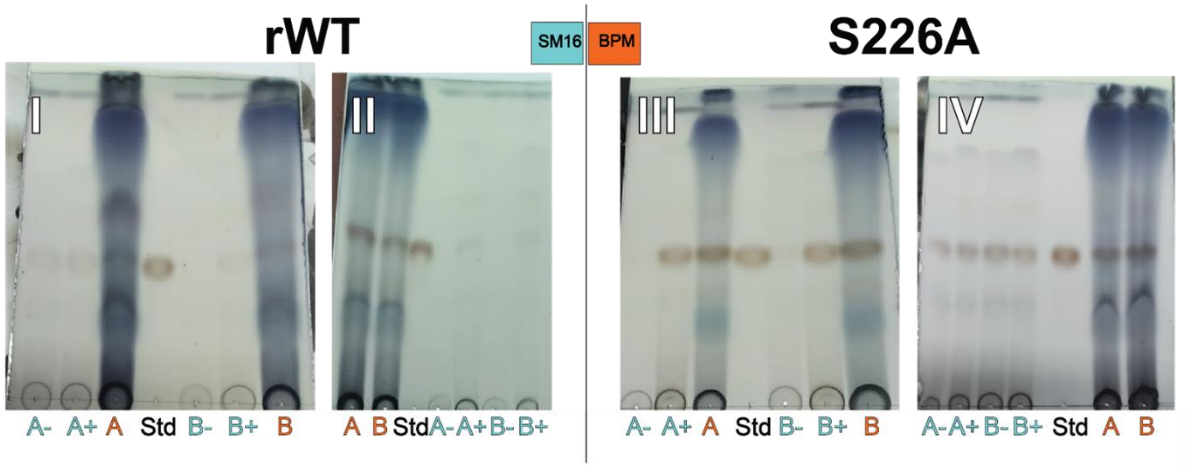


*Figure S20: Thin-layer chromatography to test the effects of medium and mutation. Each plate corresponds to one clone (I–IV) grown under different media conditions (SM16 in blue, BPM in orange). Each clone was tested with two biological replicates (A and B). The plus (+) and minus (–) signs indicate the addition or absence, respectively, of 5 g/L PEG 8000 in SM16 medium. Each spot was 20 µL; more intense or larger spots indicate higher concentration of the compound. BPM samples were reconstituted with double the volume, so their intensity represents half of the actual concentration. The line marked as Std corresponds to 87 mg/L of premonensin. The red spots at Rf ≈ 0.4 correspond to premonensin. The blue smeared spot at Rf ≈ 1 corresponds to residual oil from the extract.*

For the densiometric analyses of the TLC, all samples were spotted using 20 µL of crude fermentation extracts in MeOH. BPM samples were prepared at half the concentration, so the spots represent half the actual titers. It is very important to add an external calibration curve to each TLC plate as the vanillin reaction can differ slightly from plate to plate.

The densitometric values were adjusted with a natural logarithmic curve. This is because the signal loses linearity at the calibration range. This is different from direct measurement with vanillin in which the linearity is higher. After corrections by dilutions, the densitometric values turn out to be very reproducible, and the difference between biological replicates is quite small in most cases.

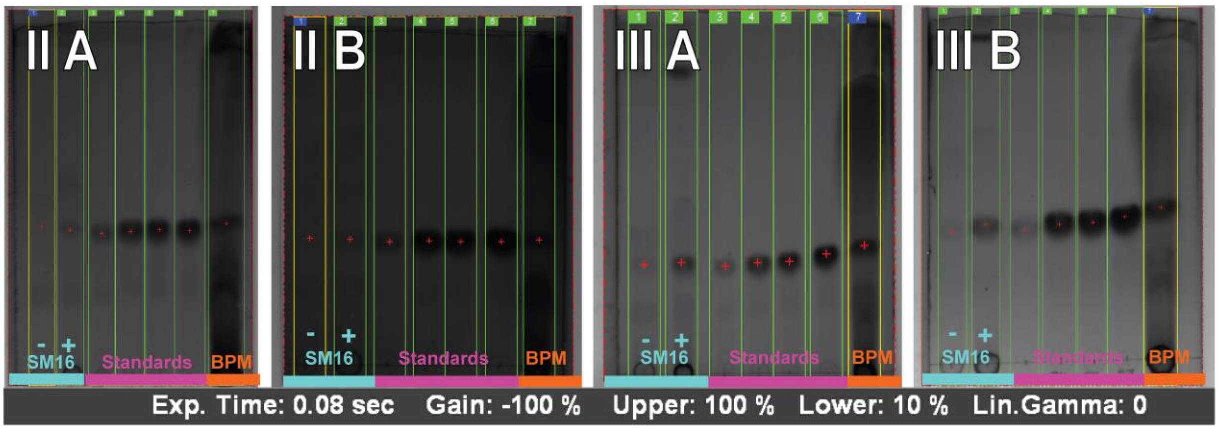


*Figure S21: Thin-layer chromatography used for the analysis of premonensin titers. Each plate corresponds to one clone, with a standards included, grown under different media conditions (SM16 in blue, BPM in orange). External premonensin standards are shown in magenta. The plus (+) and minus (–) signs indicate the presence or absence, respectively, of 5 g/L PEG 8000 in SM16 medium. Analysis lines are marked with green and yellow, premonensin spots with red crosses. Detailed processing steps are in Supplementary Information B.*

*Table S 7- Calculation of premonensin titers by densitometry and correction by dilution.*

| Sample | Densitometry (a.u. x pixel) | Concentration according to curve (mg/L) | Fermentation Volume (ml) | Reconstitution volume (ml) | Dilution | Corrected Concentration (mg/L) |
|---|---|---|---|---|---|---|
| 835 SB- | - | - | - | - | - | - |
| 835 SB+ | 4,158.00 | 19.0 | 11.5 | 5 | 0.4 | 8.2 |
| 835 BB | 10,755.00 | 27.9 | 5 | 10 | 2.0 | 55.9 |
| 835 SA- | - | - | - | - | - | - |
| 835 SA+ | 6,953.00 | 18.1 | 11.5 | 5 | 0.4 | 7.9 |
| 835 BA | 20,036.00 | 30.8 | 4 | 10 | 2.5 | 77.0 |
| 960 SA- | 5,733.00 | 21.0 | 12 | 5 | 0.4 | 8.7 |
| 960 SA+ | 34,564.00 | 54.2 | 10.5 | 5 | 0.5 | 25.8 |
| 960 BA | 35,120.00 | 55.2 | 4 | 10 | 2.5 | 138.0 |
| 960 SB- | 9,017.00 | 21.0 | 12 | 5 | 0.4 | 8.8 |
| 960 SB+ | 47,110.00 | 49.8 | 11 | 5 | 0.5 | 22.6 |
| 960 BB | 49,519.00 | 52.5 | 4 | 10 | 2.5 | 131.4 |

### I) General materials and Methods

#### 1) Materials

Glass beads (4 mm diameter) for small cultures were purchased from VWR. Compression springs, custom-made from food-grade stainless steel 18/10 ($l_0$ =250 mm, Di = 12 mm, d = 1 mm, m = 7 mm, Sodemann), were used. Petri dishes (plastic, Ø 8 cm, sterile) and Falcon tubes for precultures (12 ml, push cap, sterile) were obtained from Sarstedt. Erlenmeyer flasks (narrow neck) were sourced from VWR, and paper stoppers were from Carl Roth. Rubber stoppers for extraction (DIN 12871, natural rubber, 40 Shore A) were also purchased from VWR. The liquid culture shaker was a Multitron from Infor HT (NRS-000115524-003). Flat bottom clear polypropylene 96-well plates were purchased from Sarstedt.

#### 2) Media recipes

All quantities listed below are meant for the preparation of 1L of the respective medium. Due to previous experiences of the lab with contaminations due to the soy flour, the soy flour was autoclaved a total of two times. The first time only the soy with ultrapure water (5 cm higher than the solids) was autoclaved, later the complete medium was autoclaved. The oil was all human consumption grade bought in supermarket/online. It was kept in brown bottles or covered with aluminum foil, in the dark. Flaxseed oil was kept in the fridge when not in used (4°C). Metal magnetic covered magnetic stirrer was added before autoclaving in BGM and BPM medium and used (with stirrer plate) to homogenize the media to ensure good mixture and reproducible results. Solid media were melted in microwave.

**TSB**

| | | |
|---|---|---|
| Tryptic Soy Broth | 30 g | |
| Add $H_2O$ to | 1.0 l | autoclave |

**SM-16**

| | | |
|---|---|---|
| MOPS | 20.90 g | |
| L-Proline | 10.00 g | |
| Glucose | 20.00 g | |
| NaCl | 0.50 g | |
| $K_2HPO_4$ | 2.10 g | |
| EDTA | 0.25 g | |
| $MgSO_4\cdot(H_2O)_7$ | 0.49 g | |
| $CaCl_2\cdot(H_2O)_2$ | 0.029 g | |
| 10× Trace Element Solution | 1 ml | |
| add MQ $H_2O$ to | 1 l | adjust to pH = 7.0 (NaOH) and autoclave. |
| $MgCl_2$ (1 M, autoclaved) | 20 ml | |

**10× Trace Element Solution**

| | |
|---|---|
| $Na_2EDTA$ | 2.000 g |

$FeSO_4$ 1.800 g
$ZnSO_4$ 860 mg
$MnSO_4$ 223 mg
$CuSO_4 \cdot (H_2O)_5$ 125 mg
Borax ($NaB_4O_7 \cdot (H_2O)_7$) 95 mg
$CoCl_2 \cdot (H_2O)_6$ 48 mg
$(NH_4)_6Mo_7O_{24} \cdot (H_2O)_4$ 35 mg
add MQ $H_2O$ to 100 ml adjust to pH = 3.5 with dil. $H_2SO_4$, sterile filter.

**Bulgarian Germination Medium**

Soy flour (previously autoclaved) 15.0 g
Dextrin 20.0 g
Glucose 5.0 g
$CaCO_3$ 1.0 g
Dry yeast 1.5 g
add MQ $H_2O$ to 1.0 l adjust to pH = 6.4, autoclave.

**Bulgarian Production Medium**

Soy flour (previously autoclaved) 32.6 g
Glucose 20.0 g
Methyl oleate 10.66 ml
$Na_2SO_4$ (anhyd.) 2.2 g
$CaCO_3$ 2.0 g
$Al_2(SO_4)_3$ 705 mg
$MnCl_2$ 330 mg
Antifoam A (approx.) 200 mg
$FeSO_4$ 110 mg
$K_2HPO_4$ 75 mg
*L*-Ascorbic acid 19 mg
add MQ $H_2O$ to 1.0 l adjust to pH = 6.6 - 6.9 (NaOH), autoclave.

Autoclaved soy oil is added to each flask at the beginning (6%) and after 8 days (3%). For the rutinary quantifications, soybean oil was used. For the feeding experiment, this was replaced with the respective oil in the same volumes.

**SFM or MS agar medium**

Soy flour (previously autoclaved) 20.0 g
Mannitol 20.0 g
Agar 20.0 g
Add MQ $H_2O$ to 20.0 g 1.0 l adjust to pH = 7.0 (NaOH), autoclave.

**Gym agar**

Yeast extract 4.0 g
Glucose 4.0 g

| | | | |
|---|---|---|---|
| Malt extract | 10 g | | |
| $CaCO_3$ | 2 g | | |
| add approximately MQ water to | 0.5 l | | adjust to pH = 7.2 |
| Agar | 23 g | autoclave | |

#### 3) Fermentation procedures

##### Solid Medium

Solid cultures (25 ml) were incubated at 30°C for 7 days before collecting spores/picking single colonies/cutting plugs.

##### Preparation of stock solutions of spores

Spore stocks were prepared at room temperature, working quickly to prevent germination. A 6 mL syringe (without plunger) was placed inside a 15 mL centrifuge tube, and a cotton plug was inserted and compacted with sterile tweezers after wetting with autoclaved physiological solution (9 g/L NaCl and 4 g/L MgSO4, pH 7) to form a filter. Approximately 5 mL of physiological solution was added to the agar plate, and spores were gently scraped from the mycelium with a sterile rake to generate a dense suspension. The suspension was filtered through the cotton-packed syringe into the centrifuge tube, avoiding full plunger compression. The filtrate was mixed with sterile 80% glycerol to a final concentration of 20% (v/v), vortexed, aliquoted into sterile 1.5 mL microcentrifuge tubes and stored at -80 °C.

##### Liquid Cultures

Fermentations were initiated from TSB precultures (3 mL in 12 mL centrifuge tubes with 6–8 glass beads, 4 mm, or 15 mL in 250 mL Erlenmeyer flasks with one glass bead spoon). Inoculation methods were either 1 agar plug per 3 mL culture (monensin quantification), 1 $cm^2$ agar (premonensin and derivatives quantification) or 1 mL of frozen mycelium stock (oil additives). Precultures were grown at 30 °C, 180 rpm for 24 h (Erlenmeyer) or 48 h (centrifuge tubes). All SM16 fermentations had a total volume of 15 ml of media and were grown in 250 ml Erlenmeyer flasks inoculated with 5% (v/v) preculture and grown with glass beads and foam stoppers for 5 days. In the premonensin strains in SM16, the medium was supplemented with 20 g/L XAD16 resin.. BPM fermentations included 24 h in germination medium (with glass beads and foam stoppers) followed by 12 days of cultivation (with springs and paper stoppers). Oil was added on the first day (6% vol) and the eight day (3% volume) of the BPM fermentations. All liquid cultures were incubated at 30 °C, 180 rpm, with a 5 cm throw.

##### Extraction of premonensin and intermediates

Resin and cell paste were followingly harvested by centrifugation (25 min at 4 °C, 3900 rpm), transferred to 15 mL centrifuge tubes, and stored at –80 °C overnight. The next day, samples were thawed at RT, and 6 mL of ethyl acetate and 1 spoon of glass beads were added to a 15 mL centrifuge tube, followed by vortexing for 30 s. Then all samples were extracted overnight at 19°C, 180 rpm in a horizontal position. Afterwards samples were centrifuged for 10 min at 4°C, 3900 rpm. 4 mL of the organic phase was transferred to test tubes using and the solvent was evaporated at 38 °C. Residues were reconstituted in 1 ml LC-MS grade acetonitrile and transferred to 1.5 mL microcentrifuge tubes, and frozen at -20°C overnight. Afterwards the tubes are centrifuged for 30 min at 3900 rpm, 4°C to sediment suspended particles. Finally, the supernatants were transferred to HPLC vials and 2 µL (A495 variants)/5 µL (DH4(0) variant) was injected into LC-MS.

**Large scale fermentation for qNMR analysis**

3 cm² of agar of each clone was inoculated into 15 mL TSB medium and cultivated at 30 °C, 180 rpm for 2 days in a Multitron Standard 3-stack system. 75 mL preculture was inoculated into 1.5 L SM-16 medium supplemented with 20 g/L XAD-16 resin. Fermentation was caried out for 5 days at 30 °C, 180 rpm, after which the resin and cell paste were harvested by centrifugation (25 min at 4 °C, 3900 rpm), transferred to 50 mL centrifuge tubes, and stored at –80 °C overnight.

**Premonensin extraction for qNMR analysis**

The frozen cell and XAD resin paste was snap-frozen in liquid nitrogen, mortared, and freeze-dried (with dry weight recorded). An overnight extraction was performed at 19 °C, 180 rpm with glass beads (1.7–2.1 mm) and ethyl acetate at a ratio of 1 g sample : 5 g beads : 20 mL solvent. Two additional 1 h extractions with 300 mL ethyl acetate followed, each with centrifugation for collecting the supernantant (20 min, 4 °C, 3900 rpm). The combined organic phases were dried over magnesium sulfate and concentrated under reduced pressure at 30 °C to yield a crude extract. The residue was re-dissolved in a minimal amount of ethyl acetate, adsorbed onto silica gel, and further concentrated by rotary evaporation. The sample was then loaded onto a pre-packed column (≈35 g silica gel in a 3.5 cm column, prepared with cyclohexane) and fractionated by sequential elution with cyclohexane/ethyl acetate mixtures (500 mL 80/20, 500 mL 60/40, then 500 mL 50/50). Fractions containing premonensin (Rf value is 0.37, mobile phase: 40/60 cyclohexane/ethyl acetate), as determined by TLC with vanillin staining, were pooled and concentrated under vacuum.

**Fermentation for Monensin Production**

Tryptic soy broth (TSB) precultures were prepared with agar plugs (1 ml tip size) as inoculum. Aliquots of this preculture (5% v/v) were transferred to Bulgarian germination medium (BGM) containing soy flour and glass beads. After two days, BPM cultures were prepared (10% v/v inoculum/medium) and incubated for 12 days. For comparison, the same precultures were used to inoculate both BGM and SM16 media. Due to the shorter incubation time of the SM16 fermentation, after the five days, the cultures were transferred to 15 ml centrifuge tubes and frozen at -80° C until the extraction day. Both media were extracted and quantified at the same time.

**Monensin extraction**

Complete extraction of the BPM medium (instead of taking aliquots) turned to be the most reproducible and complete way to extract monensin since some *Streptomyces* mycelia can get stuck to the Erlenmeyer walls and cause titers' underestimations. Extraction solvent (ethyl acetate, 0.66 eq in volume) was then added directly into the

Erlenmeyer flasks and paper stoppers were replaced by rubber stoppers and the extraction was carried out in the shaker (2h, 30°C, 180 rpm). Extractions were repeated twice more (each 0.33 eq ethyl acetate), for 30´ (at 30°C, 180 rpm). Organic fractions were collected, combined and evaporated in vacuo until only oily residue remained.

**Premonensin Thin Layer Chromatography**

Figures 9 A and B were extracted from the Figures S17 and S18, respectively. For Figure 9A, the spots were just arranged for easier inspection, no color editing was applied to this.

For Figure 9B: after carrying out the TLC and coloring with the dye, it was immediately photographed using Intas camera keeping the settings and zoom the same for all TLCs (Exposure time 0.08 seconds, Gain -100%, Upper 100%, lower 10%, Lin.Gamma=0). One TLC, with premonensin external calibration curve was done per clone. Premonensin external calibration curve was prepared using Premonensin extracted as mentioned previously and quantified with qNMR as before. Pictures were cropped using Adobe Illustrator and combined together. The only adjustment was setting the opacity of both images to 95% to improve visibility and then adjusting in Powerpoint365 the brightness to 36% and the contrast to 42%.

#### 4) HPLC Analysis for premonensin and intermediates

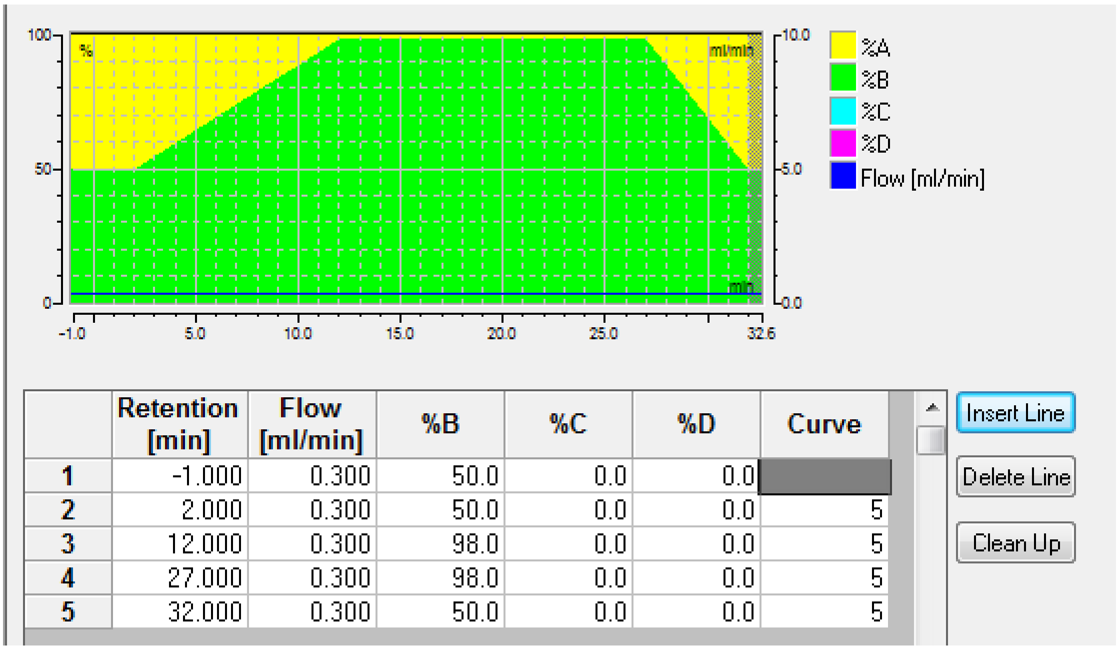


| | Retention [min] | Flow [ml/min] | %B | %C | %D | Curve |
|---|---|---|---|---|---|---|
| 1 | -1.000 | 0.300 | 50.0 | 0.0 | 0.0 | |
| 2 | 2.000 | 0.300 | 50.0 | 0.0 | 0.0 | 5 |
| 3 | 12.000 | 0.300 | 98.0 | 0.0 | 0.0 | 5 |
| 4 | 27.000 | 0.300 | 98.0 | 0.0 | 0.0 | 5 |
| 5 | 32.000 | 0.300 | 50.0 | 0.0 | 0.0 | 5 |

*Figure S22: HPLC* method used for extracts measurement. For internal calibration a lock mass of 622.028960 (Hexakis(1H,1H,2H-perfluoroethoxy)phosphazene) and sodium formate clusters were used.

#### MS Method

##### Method Set: D:\Methods\Susanna\110-1300 autoMSMS pos_.m

BRUKER

### otofControl

#### General Information

| | | | |
|---|---|---|---|
| **Method Name:** | 110-1300 autoMSMS pos_.m | **Saved:** | 2018/09/05 09:38:08+02:00 |
| **Application Name:** | Bruker otofControl | **Application Version:** | 4.1.3.5 |
| **Device Type:** | compact | **Device Serial Number:** | 8255754.20147 |
| **Operator:** | Demo User | **Host:** | COMPACT-20147 |
| **Operating System:** | Windows 7 Professional | **Organisation:** | Bruker Daltonik GmbH |

#### Chromatogram

##### Chromatogram Traces

| Enabled | Color | Type | Masses | Width | Polarity | Filter |
|---|---|---|---|---|---|---|
| On | Red | BPC | | | ± | MS |
| On | Blue | TIC | | | ± | MS |
| On | Black | TIC | | | ± | All MS/MS |

#### SPL

| | |
|---|---|
| **Scheduled Precursor List:** | Off |

#### Segment 1

#### 0 .... 0.02 min

##### Main

| | | | |
|---|---|---|---|
| **Polarity:** | Positive | **Scan Mode:** | MS |
| **Mass Range from:** | 110 m/z | **Mass Range to:** | 1300 m/z |
| **Rolling Average:** | Off | **Rolling Average No.:** | 2 |
| **Spectra rate:** | 8.00 Hz | **View:** | Expert |

##### Mode

| | | | |
|---|---|---|---|
| **Save Spectra:** | Line and Profile Spectra | **Line Spectra Calculation:** | Use Maximum Intensity |
| **Absolute Threshold (per 1000 sum.):** | 25 cts. | **Peak Summation Width:** | 3 pts. |
| | | **Focus Active:** | Off |
| **Mark as Calibration Segment:** | Off | | |

##### Source

| | | | |
|---|---|---|---|
| **Source:** | ESI | | |
| **End Plate Offset:** | 500 V | **Capillary:** | 4500 V |
| **Nebulizer:** | 2.2 Bar | **Dry Gas:** | 10.0 l/min |
| **Dry Temp:** | 220 °C | **Divert Valve:** | Waste 1-6 |

##### Tune

| | | | |
|---|---|---|---|
| **Funnel 1 RF:** | 150.0 Vpp | **Funnel 2 RF:** | 200.0 Vpp |
| **isCID Energy:** | 0.0 eV | **Hexapole RF:** | 50.0 Vpp |
| **Ion Energy:** | 4.0 eV | **Low Mass:** | 90.0 m/z |
| **Collision Energy:** | 7.0 eV | **Pre Pulse Storage:** | 5.0 µs |
| **Stepping:** | On | **Mode:** | Basic |
| **Collision RF from:** | 550.0 Vpp | **Collision RF to:** | 550.0 Vpp |
| **Transfer Time from:** | 80.0 µs | **Transfer Time to:** | 80.0 µs |
| **Timing from:** | 50 % | **Timing to:** | 50 % |
| **Collision Energy from:** | 100 % | **Collision Energy to:** | 250 % |
| **Timing from:** | 50 % | **Timing to:** | 50 % |

### MS/MS

**Auto MS/MS:** Off

##### *MRM*

**MRM:** Off

##### *isCID*

**isCID (MS-MS/MS):** Off

##### *bbCID*

**bbCID (MS-MS/MS):** Off

#### Segment 2

#### 0.02 .... 0.3 min

##### Main

| | | | |
|---|---|---|---|
| **Polarity:** | Positive | **Scan Mode:** | MS |
| **Mass Range from:** | 110 m/z | **Mass Range to:** | 1300 m/z |
| **Rolling Average:** | Off | **Rolling Average No.:** | 2 |
| **Spectra rate:** | 8.00 Hz | **View:** | Expert |

##### Mode

| | | | |
|---|---|---|---|
| **Save Spectra:** | Line and Profile Spectra | **Line Spectra Calculation:** | Use Maximum Intensity |
| **Absolute Threshold (per 1000 sum.):** | 25 cts. | **Peak Summation Width:** | 3 pts. |
| | | **Focus Active:** | Off |
| **Mark as Calibration Segment:** | On | | |

##### Source

| | | | |
|---|---|---|---|
| **Source:** | ESI | | |
| **End Plate Offset:** | 500 V | **Capillary:** | 4500 V |
| **Nebulizer:** | 2.2 Bar | **Dry Gas:** | 10.0 l/min |
| **Dry Temp:** | 220 °C | **Divert Valve:** | Source 1-2 |

##### Tune

| | | | |
|---|---|---|---|
| **Funnel 1 RF:** | 150.0 Vpp | **Funnel 2 RF:** | 200.0 Vpp |
| **isCID Energy:** | 0.0 eV | **Hexapole RF:** | 50.0 Vpp |
| **Ion Energy:** | 4.0 eV | **Low Mass:** | 90.0 m/z |
| **Collision Energy:** | 7.0 eV | **Pre Pulse Storage:** | 5.0 µs |
| **Stepping:** | On | **Mode:** | Basic |
| **Collision RF from:** | 550.0 Vpp | **Collision RF to:** | 550.0 Vpp |
| **Transfer Time from:** | 80.0 µs | **Transfer Time to:** | 80.0 µs |
| **Timing from:** | 50 % | **Timing to:** | 50 % |
| **Collision Energy from:** | 100 % | **Collision Energy to:** | 250 % |
| **Timing from:** | 50 % | **Timing to:** | 50 % |

### MS/MS

| | |
|---|---|
| **Auto MS/MS:** | Off |

##### *MRM*

| | |
|---|---|
| **MRM:** | Off |

##### *isCID*

| | |
|---|---|
| **isCID (MS-MS/MS):** | Off |

##### *bbCID*

| | |
|---|---|
| **bbCID (MS-MS/MS):** | Off |

#### Segment 3
#### 0.3 .... unlimited min

##### Main

| | | | |
|---|---|---|---|
| **Polarity:** | Positive | **Scan Mode:** | Auto MS/MS |
| **Mass Range from:** | 110 m/z | **Mass Range to:** | 1300 m/z |
| **Rolling Average:** | Off | **Rolling Average No.:** | 2 |
| **Spectra rate:** | 8.00 Hz | **View:** | Expert |

##### Mode

| | | | |
|---|---|---|---|
| **Save Spectra:** | Line and Profile Spectra | **Line Spectra Calculation:** | Use Maximum Intensity |
| **Absolute Threshold (per 1000 sum.):** | 25 cts. | **Peak Summation Width:** | 3 pts. |
| | | **Focus Active:** | Off |
| **Mark as Calibration Segment:** | Off | | |

##### Source

| | | | |
|---|---|---|---|
| **Source:** | ESI | | |
| **End Plate Offset:** | 500 V | **Capillary:** | 4500 V |
| **Nebulizer:** | 2.2 Bar | **Dry Gas:** | 10.0 l/min |
| **Dry Temp:** | 220 °C | **Divert Valve:** | Waste 1-6 |

#### Tune

| | | | |
|---|---|---|---|
| **Funnel 1 RF:** | 150.0 Vpp | **Funnel 2 RF:** | 200.0 Vpp |
| **isCID Energy:** | 0.0 eV | **Hexapole RF:** | 50.0 Vpp |
| **Ion Energy:** | 4.0 eV | **Low Mass:** | 90.0 m/z |
| **Collision Energy:** | 7.0 eV | **Pre Pulse Storage:** | 5.0 µs |
| **Stepping:** | On | **Mode:** | Basic |
| **Collision RF from:** | 550.0 Vpp | **Collision RF to:** | 550.0 Vpp |
| **Transfer Time from:** | 80.0 µs | **Transfer Time to:** | 80.0 µs |
| **Timing from:** | 50 % | **Timing to:** | 50 % |
| **Collision Energy from:** | 100 % | **Collision Energy to:** | 250 % |
| **Timing from:** | 50 % | **Timing to:** | 50 % |

## MS/MS

| | | | |
|---|---|---|---|
| **Auto MS/MS:** | On | | |
| **Precursor Ion List:** | Exclude | **Cycle Time:** | 0.5 sec |
| **Threshold (per 1000 sum.) Absolute:** | 400 cts | **Active Exclusion:** | On |
| | | **Exclude after:** | 3 Spectra |
| | | **Reconsider Precursor:** | On |
| **Release after:** | 0.20 min. | **Smart Exclusion:** | Off |
| **if Curent Intens./Prev. Intens.:** | 1.8 | | |
| **Smart Exclusion:** | 2 x | | |

##### Exclude Mass List

| Mass Range Start | Mass Range End | |
|---|---|---|
| 102.08 | 102.18 | 1 |
| 621.98 | 622.08 | 2 |
| 643.96 | 644.06 | 3 |
| 659.94 | 660.04 | 4 |

##### Auto MS/MS Preference

| | | | |
|---|---|---|---|
| **Preferred Range:** | Off | | |
| **Preferred Range Low:** | 2 | **Preferred Range High:** | 5 |
| **Exclude Singly:** | Off | **Exclude unknown:** | Off |
| **Group Length:** | 3 | **Strict Active Exclusion:** | Off |
| **Sort Precursors by:** | Intensity | **Preferred mass list:** | Empty list |

##### Auto MS/MS Multi CE

**Auto MS/MS Multi CE:** Off

##### SILE

**SILE:** Off

##### CID

**Fallback Charge State:** 1 z

###### Isolation + Fragmentation List

| Type | Mass [m/z] | Width [m/z] | Collision Energy [eV] | Charge State | |
|---|---|---|---|---|---|
| Base | 100.00 | 4.00 | 20.0 | 1 | 1 |
| Base | 500.00 | 5.00 | 20.0 | 1 | 2 |
| Base | 1000.00 | 6.00 | 20.0 | 1 | 3 |
| Base | 1300.00 | 8.00 | 30.0 | 1 | 4 |

##### CID Acquisition

| | | | |
|---|---|---|---|
| **Acquisition:** | On | **Spectra Rate MS:** | 8.00 Hz |
| **MS/MS low (per 1000 sum.):** | 10000.0 cts. | **MS/MS low:** | 1600 x |
| **MS/MS high:** | 100000.0 cts. | **MS/MS high:** | 800 x |
| **Total Cycle Time Range:** | n/a sec | **Absolute Threshold :** | n/a cts. |

##### MRM

**MRM:** Off

##### isCID

**isCID (MS-MS/MS):** Off

##### bbCID

**bbCID (MS-MS/MS):** Off

### Publication bibliography


Bierman, M.; Logan, R.; O'Brien, K.; Seno, E. T.; Rao, R. N.; Schoner, B. E. (1992): Plasmid cloning vectors for the conjugal transfer of DNA from Escherichia coli to Streptomyces spp. In *Gene* 116 (1), pp. 43–49. DOI: 10.1016/0378-1119(92)90627-2.

Breaud, Célia; Lallemand, Laura; Mares, Gary; Mabrouki, Fathi; Bertolotti, Myriam; Simmler, Charlotte et al. (2022): LC-MS Based Phytochemical Profiling towards the Identification of Antioxidant Markers in Some Endemic Aloe Species from Mascarene Islands. In *Antioxidants (Basel, Switzerland)* 12 (1). DOI: 10.3390/antiox12010050.

Chambers, Matthew C.; Maclean, Brendan; Burke, Robert; Amodei, Dario; Ruderman, Daniel L.; Neumann, Steffen et al. (2012): A cross-platform toolkit for mass spectrometry and proteomics. In *Nature Biotechnology* 30 (10), pp. 918–920. DOI: 10.1038/nbt.2377.

Edgar, Robert C. (2004): MUSCLE: a multiple sequence alignment method with reduced time and space complexity. In *BMC bioinformatics* 5, p. 113. DOI: 10.1186/1471-2105-5-113.

Kushnir, Susanna; Sundermann, Uschi; Yahiaoui, Samir; Brockmeyer, Andreas; Janning, Petra; Schulz, Frank (2012): Minimally Invasive Mutagenesis Gives Rise to a Biosynthetic Polyketide Library. In *Angew. Chem. Int. Ed.* 51 (42), pp. 10664–10669. DOI: 10.1002/anie.201202438.

Zdouc, Mitja M.; Blin, Kai; Louwen, Nico L. L.; Navarro, Jorge; Loureiro, Catarina; Bader, Chantal D. et al. (2025): MIBiG 4.0: advancing biosynthetic gene cluster curation through global collaboration. In *Nucleic Acids Res* 53 (D1), D678-D690. DOI: 10.1093/nar/gkae1115.
